# Asymmetric crosstalk between the BMP and TGFβ pathways resolves signaling ambiguity

**DOI:** 10.64898/2026.04.20.719570

**Authors:** Johannes M. Auth, Inbal Eizenberg-Magar, Omer Erez, Omer Zachar, Merav D. Shmueli, Inbal Zigdon, Achinoam Shoham, Vladimir Mindel, Meytar Asulin Berrebi, Ariel Tennenhouse, Neta Bar-Hai, Rakefet Ben-Yishay, Dana Ishay-Ronen, Yaron E. Antebi

**Affiliations:** Department of Molecular Genetics, Weizmann Institute of Science, Rehovot, Israel; Department of Biomolecular Sciences, Weizmann Institute of Science, Rehovot, Israel; Department of Systems Immunology, Weizmann Institute of Science, Rehovot, Israel; Oncology Institute, Sheba Medical Center, Ramat-Gan, Israel; Faculty of Medicine, Tel Aviv University, Tel Aviv, Israel

## Abstract

The BMP and TGFβ signaling pathways control cellular fate decisions in diverse biological contexts, often playing opposing roles. Despite extensive knowledge of these pathways, understanding how cells respond to environments containing these opposing cues remains a challenge. Here, we systematically analyze the activation of these pathways under combinatorial signaling environments. We find that TGFβ ligands inhibit BMP signaling, while BMP ligands enhance TGFβ transcriptional response across concentrations, ligand variants, and cell types. This asymmetric crosstalk results in the activation of a TGFβ-biased transcriptional response, even under mixed signaling conditions, effectively reducing signal ambiguity, with implications for processes such as EMT. We show that this crosstalk originates downstream of the SMAD proteins phosphorylation. Using mathematical models, we predict, and experimentally verify, that promiscuous interactions between SMAD proteins provide the mechanism for the observed crosstalk. Our findings challenge the canonical models, suggesting an active role for mediator proteins in determining biological responses.

## Introduction

The BMP and TGFβ pathways play crucial roles in a wide range of biological processes, from early embryonic development to pathological conditions such as cancer^1–3^. Extensive research has explored the individual impacts of these pathways on specific differentiation processes. Frequently, these two pathways exert opposing effects on cellular processes. However, despite their contrasting roles, in many biological contexts, cells are simultaneously exposed to ligands from both the BMP and TGFβ families^4–6^. Consequently, these pathways provide a compelling model system for studying the combinatorial effects of opposing ligands on cellular behavior.

One pivotal process where this interplay is particularly pronounced is epithelial-to-mesenchymal transition (EMT), a process tightly regulated by BMP and TGFβ^7,8^. During EMT, epithelial cells lose their polarity, detach from the tissue, and gain motility. While EMT is essential for early embryonic development and wound healing, its dysregulation contributes to cancer progression and metastasis.

Notably, TGFβ is a predominant driver of EMT^9,10^, while BMP signaling triggers the reverse process of mesenchymal-to-epithelial transition (MET)^11,12^. Together, these pathways regulate various developmental EMT processes, such as germ layer determination during gastrulation^13–15^, and are significantly implicated in cancer pathogenesis^16^. Furthermore, these pathways exert opposing roles in mesenchymal cell differentiation^17–21^, cancer^22,23^, immune responses^24^, and various other biological processes^25^. Therefore, understanding the combinatorial effects of these signals is crucial for achieving predictive insights and control over EMT and other biological processes (Figure 1A).

**Figure 1.**
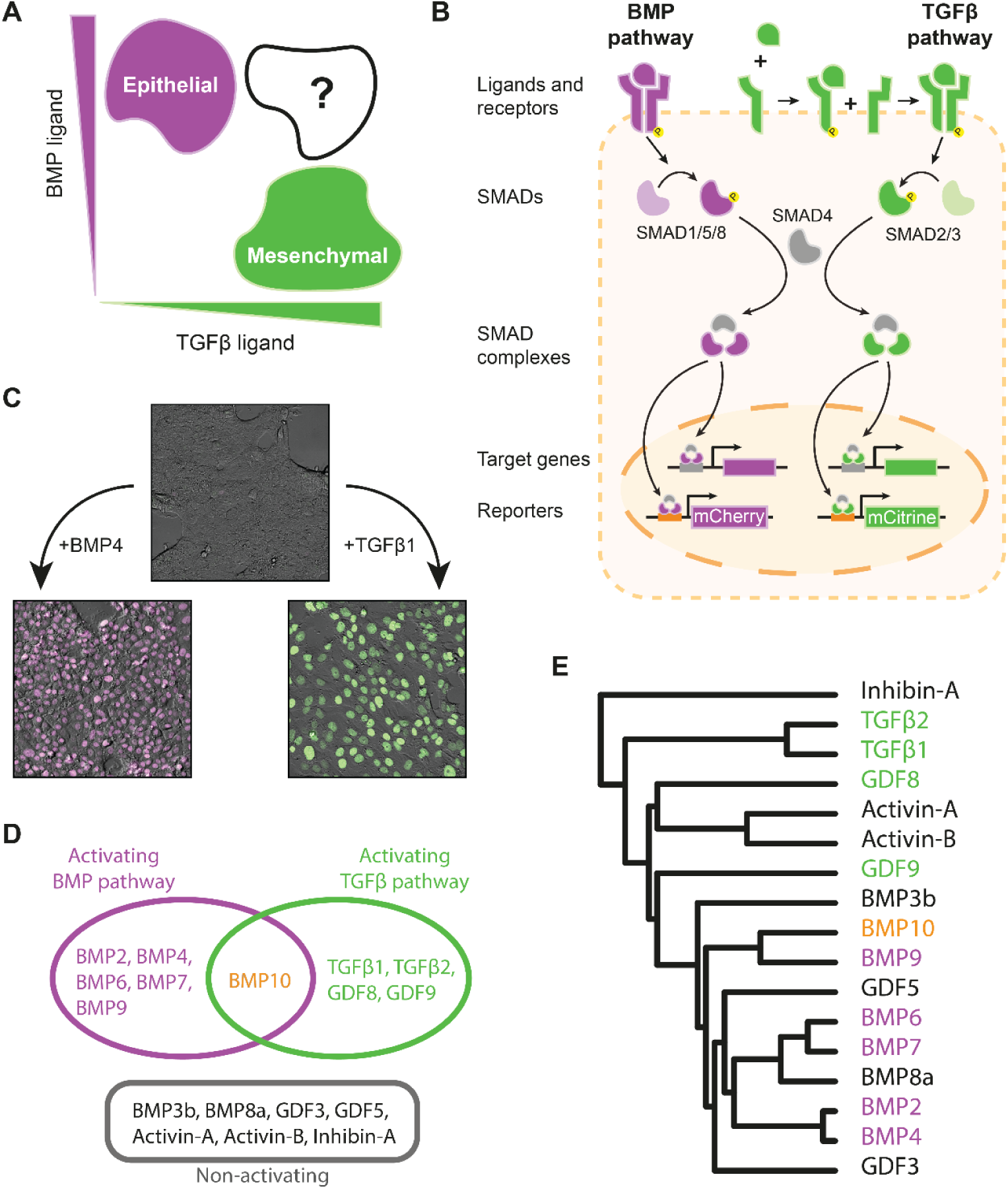
Fluorescent reporter system reveals the activation patterns of BMP and TGFβ ligands. (A) Schematic depiction of the mapping from ligand environment to the resulting cell fate in EMT. For high BMP and low TGFβ concentrations (top left), cells show an epithelial state. For low BMP and high TGFβ concentrations (bottom right), cells transition to a mesenchymal state. In multi-ligand environments (top right), cells are exposed to mixed signals, and the resulting fate depends on how this ambiguity is resolved. (B) Schematic depiction of the molecular components within the BMP and TGFβ pathways (purple and green, respectively). The binding of ligands and receptors induces phosphorylation and complex formation of SMAD family proteins, which act as transcription factors. The activity level of each pathway can be quantified by introducing a pathway-specific transcriptional response element controlling the expression of a fluorescent reporter gene (mCherry for BMP and mCitrine for TGFβ). (C) Microscopic images of the dual reporter cells after 24h of incubation with no ligand exposure (top), with either 31.25 ng/ml of BMP4 or 1.92 ng/ml of TGFβ. The mCherry channel is depicted in purple, and the mCitrine channel is depicted in green. (D) Ligands are classified based on their activation profile - BMP activating ligands (purple), TGFβ activating ligands (green), ligands activating both pathways (orange), and non-activating ligands (black). (E) Phylogenetic tree of BMP and TGFβ ligands (adapted from Chang et al.^38^). Ligands are color-coded according to their classification (cf. D). See also Figure S1.

At the molecular level, the BMP and TGFβ pathways share a similar structure that has been extensively elucidated^26–29^ (Figure 1B). Secreted ligands interact promiscuously with several receptor variants, forming active signaling complexes. Subsequently, these signaling complexes phosphorylate intracellular, pathway-specific SMAD proteins. BMP ligands activate SMAD1, SMAD5, and SMAD8, while TGFβ ligands activate SMAD2 and SMAD3. Once activated, two SMADs combine with the shared SMAD4 protein to form trimeric complexes, which then translocate to the nucleus, acting as transcription factors for the respective target genes. A noteworthy feature of the BMP and TGFβ pathways is the potential competition stemming from shared SMAD4 binding and the promiscuous interactions among ligands, receptors, and SMADs from both pathways^30–34^. Recent studies have revealed that such promiscuous interactions within the BMP pathway can bestow complex computational capabilities^35–37^. Similarly, the multi-layered promiscuous networks characterizing BMP and TGFβ pathways might provide the computational capacity required to resolve ambiguity between the corresponding signals.

In this study, we examine how cells perceive signals within complex signaling environments and identify the role of SMAD interactions in resolving ambiguous signals. We find that the BMP and TGFβ pathways activate independently in response to their respective ligands. However, when cells encounter a multi-ligand environment encompassing both BMP and TGFβ-related ligands, an unexpected combinatorial crosstalk emerges. This observed crosstalk displays a bias towards the activation of the TGFβ pathway over the BMP pathway, yielding distinct regimes where either the BMP or TGFβ pathway predominates. Consequently, cells exhibit reduced transcriptional response ambiguity even under complex signaling conditions. To unravel these observations, we have constructed a mathematical model that delineates intracellular interactions between the BMP and TGFβ pathways. Based on this model, we conclude that the observed crosstalk arises from the formation of mixed SMAD complexes involving SMADs associated with both pathways. These promiscuous interactions challenge the existing paradigm of SMAD complex formation and signal transduction in the BMP and TGFβ pathways. Furthermore, the role of these pathways in resolving combinatorial signaling environments suggests that cell-fate decisions can be determined upstream of the gene regulatory networks. Our findings reveal that rather than serving as passive signal transduction channels, networks of signaling pathways have an active role in determining cell responses.

## Results

### BMP and TGFβ ligands can be classified according to the transcriptional activation patterns of the two pathways

In order to quantitatively measure the canonical, SMAD-dependent, BMP and TGFβ pathways activity, we utilized a transcriptional reporter system. We placed a response element containing multimerized binding sites specific to the SMAD mediators of each pathway upstream of a fluorescent protein (Figure 1B). The BMP reporter contains a SMAD1/5/8 response element (BRE)^39^ controlling the expression of mCherry, while the TGFβ reporter contains a SMAD2/3 response element (TRE)^40–42^ controlling the expression of mCitrine. Using these constructs, we established a dual reporter cell line based on the NAMRU Mouse Mammary Gland (NMuMG) epithelial cell line. NMuMG cells robustly respond to both BMP and TGFβ signals and are routinely used as a model system to study EMT dynamics^9,34,43,44^. We exposed cells to either BMP4 or TGFβ1 and measured a specific induction of the BRE or TRE reporter, respectively (Figure 1C). The resulting cell line enables specific and simultaneous quantification of the transcriptional activity of each individual pathway within the same cell.

Using the cell-based system, we determined the activation profile of multiple ligands associated with the BMP and TGFβ pathways. We exposed cells to 17 ligands, each administered at several concentrations across its entire dynamic range. Using flow cytometry, we measured the total fluorescence accumulated after 24 hours, representing the total transcriptional activity of the pathway (Figure S1). Based on this data, we extracted the key biochemical parameters for each ligand by fitting to a Hill function, including the EC_50_ concentration and the saturation level (Table S1). We found that ligands can be divided into four groups based on their ability to activate BMP and/or TGFβ responses (Figure 1D). Specifically, BMP2, BMP4, BMP6, BMP7, and BMP9 exclusively signal through the BMP pathway, while TGFβ1, TGFβ2, GDF8, and GDF9 exclusively signal through the TGFβ pathway. In contrast, BMP10 induces the response of both the BMP and the TGFβ reporters within the same cell. Finally, BMP3b, BMP8a, GDF3, GDF5, Activin-A, and Inhibin-A do not induce any response. It is important to note that the response groups reflect only the canonical activation capacity of these ligands and do not eliminate a non-canonical, SMAD-independent effect. We note that at high concentrations, TGFβ1 and TGFβ2 exhibit a decreasing response. However, to focus on the crosstalk between the two pathways, we will only utilize these ligands across the lower concentration range in which the cellular response increases with ligand concentration.

We compared the classification into response groups to the sequence-based phylogenetic classification of the ligands^38^ (Figure 1E). Both classifications show a similar structure for the single activating ligands, where ligands cluster according to their associated pathway. However, while BMP10 activates both pathways, the highly homologous BMP9 only activates the BMP pathway. Non-activating ligands do not show sequence-based similarity between themselves and are clustered within the activating ligands.

### BMP and TGFβ pathways exhibit an asymmetric crosstalk across ligand variants and cell lines

To examine the crosstalk between the two pathways, we considered BMP4 and TGFβ1, each exclusively activating its corresponding pathway. We found that the BMP response is inhibited when cells are exposed to both ligands, compared to when exposed to BMP4 only (Figures 2A, S2A,B). In contrast, the response of the TGFβ pathway to stimulation by the mixture of BMP4 and TGFβ1 was increased beyond the response induced by TGFβ1 alone. Several negative regulatory mechanisms are known to exist between these pathways^22,45–48^, such as competition on receptors or SMAD4, linker region phosphorylation of SMADs, and involvement of inhibitory SMADs. While the observed negative effect of TGFβ on the BMP response is consistent with such known regulatory mechanisms, these mechanisms cannot explain the enhancement of TGFβ response by BMP4. To test the generality of the observed phenomenon in other cellular contexts, we examined the effect of the two ligands and their mixture on the chondrogenic cell line ATDC5 transfected with the BRE and TRE reporters, as in NMuMG (Figure 2B). Consistent with our observations from NMuMG cells, we find that the BMP response is inhibited under mixed ligand conditions while the TGFβ response is increased.

**Figure 2.**
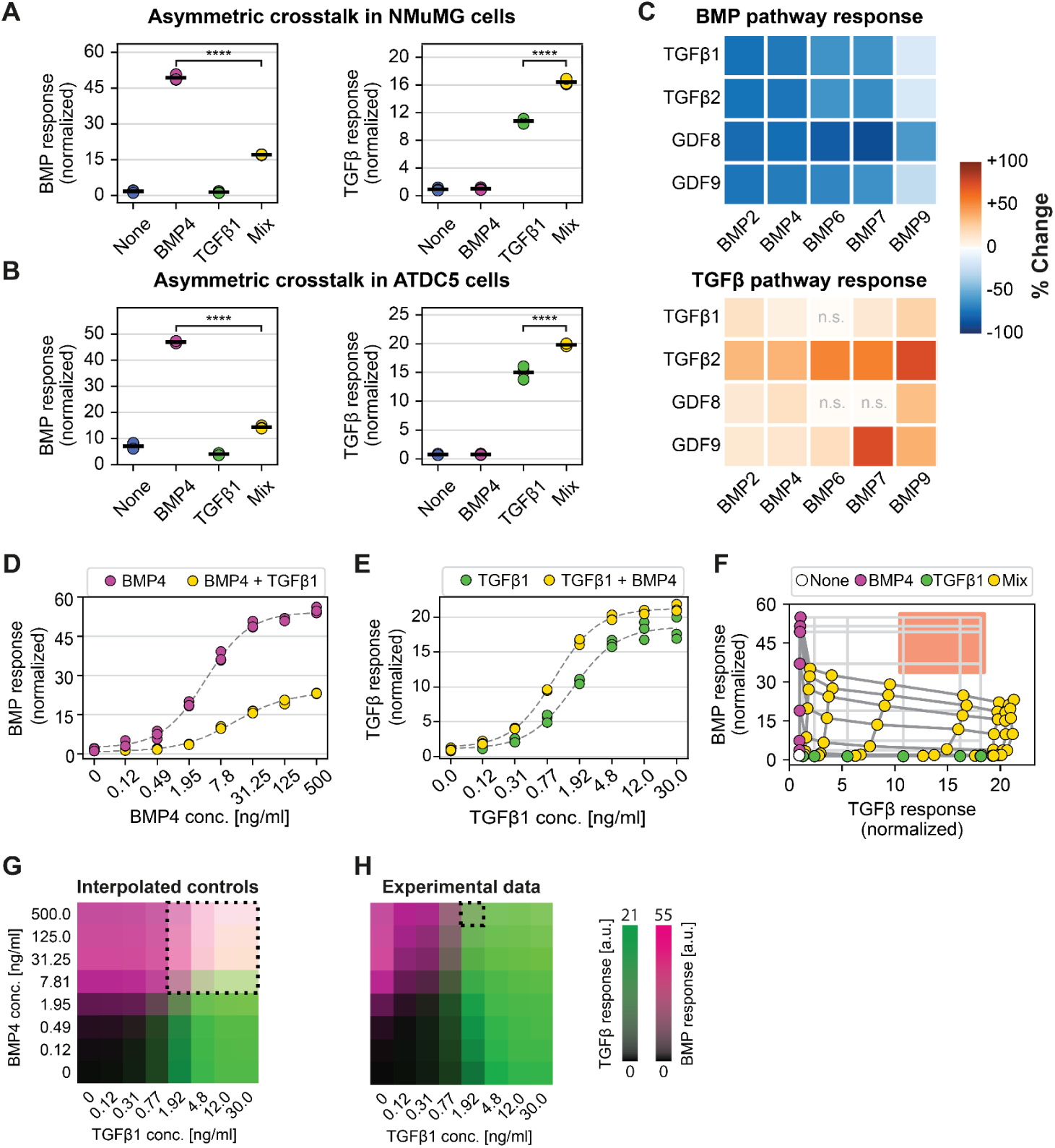
Pairwise mixed ligand conditions reveal an asymmetric crosstalk between BMP and TGFβ. (A-B) NMuMG dual reporter (A) and ATDC5 (B) cells were incubated for 24h under four different conditions: no ligands added (None), BMP4 at 31.25 ng/ml (BMP4), TGFβ1 at 1.92 ng/ml (TGFβ1), and a combination of BMP4 and TGFβ1 at these concentrations (Mix). The data shows n=3 biological repeats. (C) NMuMG cells were exposed for 24h to combinations of all pairs of a BMP-activating ligand (x-axis) with a TGFβ-activating ligand (y-axis). The percent response change was calculated and plotted for the BMP pathway (top) and TGFβ pathway (bottom) as the change between the mixed and single ligand response relative to the single ligand response (see Methods). Conditions that show no significant change in the response are marked with ‘n.s.’. The data contains n=4 biological repeats. (D) NMuMG dual reporter cells were exposed for 24h to a concentration gradient of BMP4 with (yellow) and without (purple) TGFβ1 at 12 ng/ml. Normalized fluorescence of the BMP reporter is plotted. (E) Cells were exposed for 24h to a concentration gradient of TGFβ1 with (yellow) and without (green) BMP4 at 500 ng/ml. Normalized fluorescence of the TGFβ reporter is plotted. Dashed line is a fit to a four-parameter Hill function. (C-E) Cells were exposed to two-dimensional concentration gradients using the 8×8 concentration matrix of BMP4 and TGFβ1. (F) The BMP pathway response is plotted against the TGFβ1 pathway response across the full dataset. Conditions are color-coded: no ligand (white), BMP4 only (purple), TGFβ1 only (green), and mixed conditions (yellow). Nodes of the light gray grid show the expected response coordinates of a mixed sample, assuming no crosstalk. The ambiguous region where activity levels of both pathways are over 50% of their saturation level is marked in red. (G-H) BMP and TGFβ transcriptional responses are plotted as heatmaps as a function of the BMP4 and TGFβ1 concentrations. The BMP response is shown in purple, and the TGFβ response is shown in green. White regions emerge when both pathways are activated together. Plots show interpolation of the single ligand dose responses to reflect the expected response without the crosstalk effect (G) and the actual response of the cells (H). The region where both pathways are active at levels higher than 45% of the maximal response is marked by a dashed line. In all figures, *P ≤ 0.05, **P ≤ 0.01, ***P ≤ 0.001, ****P ≤ 0.0001, determined using a two-sided t-test. See also Figures S2 and S3.

Both BMP and TGFβ pathways comprise multiple ligands. Therefore, we wanted to examine whether the asymmetric crosstalk is specific to BMP4 and TGFβ1 or whether it is a more general characteristic of the pathways. Thus, we extended our analysis to include all 20 possible pairs of a BMP-activating ligand with a TGFβ-activating ligand and measured the change in the transcriptional response (Figure 2C, S2C,D). When TGFβ ligands were added to the BMP ligands, the BMP pathway activity showed a striking reduction of up to 92% (Figures 2C, top; and S2E). In contrast, ligand pairs exhibited an enhanced TGFβ response, exceeding that of the individual TGFβ ligand (Figures 2C, bottom; and S2F). It is important to note that none of the BMP ligands activate the TGFβ pathway when administered on their own and only have an effect when added in combination with a TGFβ ligand. This data demonstrates the generality of the asymmetric combinatorial crosstalk across BMP and TGFβ ligand variants.

### BMP-TGFβ crosstalk reduces the ambiguity of signals in a TGFβ-biased manner

Since cells are exposed to different concentrations of BMP and TGFβ in their environment, we extended our measurements to determine the full concentration dependence of the asymmetric crosstalk. We started by measuring the full dose-response curve for either BMP4 or TGFβ1, both in the presence or absence of a fixed concentration of the opposing ligand. Our results show a substantial reduction of the BMP pathway response to BMP4 when administered concurrently with TGFβ1 across the entire dynamic range (Figure 2D). Conversely, the TGFβ pathway’s response to TGFβ1 was consistently enhanced in the presence of BMP4 across all concentrations (Figure 2E). Indeed, these findings were consistent across additional clones of dual-reporter cells (Figure S3A,B). To further quantify the effect of each ligand on the saturation level and EC_50_ concentration, we fitted a four-parameter Hill function to each dose response. We find that TGFβ1 decreases the saturation level of BMP4 and increases the EC_50_ concentration (Figure S3C, top). Both have a negative effect on the response of the BMP pathway. In contrast, BMP4 shows the opposite effects on the parameters of TGFβ1 (Figure S3C, bottom). Specifically, we find that the presence of BMP4 augmented the saturation level of the TGFβ pathway response, surpassing the saturation level observed in the absence of BMP4.

Next, we extended the dose-response curves to a full two-dimensional concentration matrix of BMP and TGFβ ligand pairs. All possible combinations of BMP4 and TGFβ1 concentrations were used, with each ligand administered at eight concentrations, spanning its entire dynamic range (Figure S3D), and the response of the pathways was measured. By fitting the concentration dependence of the BMP response for each TGFβ1 level, we see a gradual change in the saturation and EC_50_ (Figure S3E, top). An opposing effect is found in the TGFβ response for different BMP4 concentrations (Figure S3E, bottom). The crosstalk can be further analyzed by plotting the BMP vs TGFβ responses for each measurement (Figure 2F). The response to BMP4 only (purple) or TGFβ1 only (green) falls along the corresponding axis, suggesting an orthogonal activation of each pathway by its respective ligand. In a null model with no crosstalk between the two pathways, the response to mixed ligand conditions directly reflects the response to each ligand individually (gray orthogonal grid). In contrast, we observe a significant deviation from the no-crosstalk scenario, as the asymmetric crosstalk results in a deformation of the grid (yellow). Specifically, the BMP pathway exhibited a significantly reduced response when BMP4 was added together with TGFβ1, while the same combination yielded an elevation in the TGFβ pathway response across all concentrations. These findings were consistently observed across various ligand pairs and cell lines (Figures S3F-H).

Considering two independent pathways, a high concentration of both ligands would result in a robust activity of both pathways (Figure 2G, dotted region). In this scenario, activation of multiple distinct downstream programs will result in an ambiguous cellular response. However, studying the effect of crosstalk on the response space revealed that responses are distinctively shifted away from the region where both pathways would be strongly active simultaneously (Figure 2F, red). In fact, the asymmetric crosstalk partitions the response space into distinct regions, with each region exclusively characterized by the activation of either the BMP pathway or the TGFβ pathway (Figure 2H). Thus, the crosstalk reduces ambiguity at the cellular response level even under complex extracellular environments.

### BMP enhances TGFβ-induced transcriptional response and the Epithelial to Mesenchymal Transition program

The reporter system we use measures the transcriptional activity of the SMAD mediators on exogenous synthetic targets accumulated over 24 hours. To assess whether the asymmetric crosstalk extends to endogenous targets, we directly measured gene expression levels. To capture early responses that precede transcriptional feedback, we performed RNA sequencing on NMuMG cells cultured for 6 hours under four conditions: no ligands added, BMP4 only, TGFβ1 only, and the mix of both. To isolate the crosstalk effect, we focused on genes that respond only to one signal, either BMP4 (*Id1*, *Id2*, *Id3*) or TGFβ1 (*Serpine1*, *Snai1*, *Mmp9*), and examined their expression levels. All three BMP4 target genes were significantly suppressed under mixed conditions compared with their response to BMP4 alone (Figure 3A), and in agreement with our reporter-based observations. Similarly, the TGFβ target genes reflected the asymmetric crosstalk as they were significantly elevated in the mixed sample compared to the TGFβ1 only sample (Figure 3B). These results demonstrate that the asymmetric crosstalk is evident at the RNA level, emerges at early timepoints, and affects endogenous downstream target genes of the two pathways.

**Figure 3.**
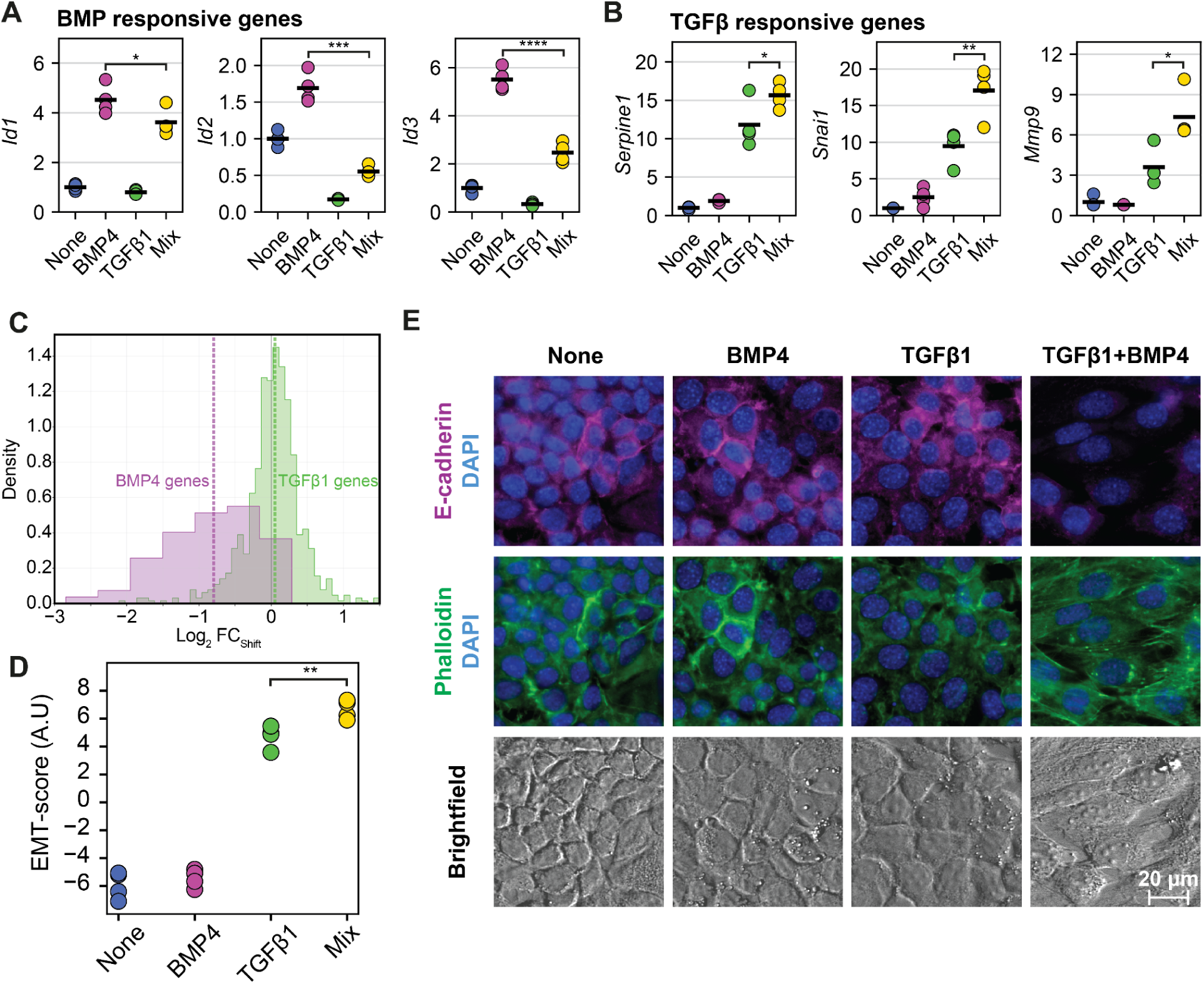
Asymmetric Crosstalk is reflected in the transcriptional level and enhances TGFβ-induced EMT. (A-C) NMuMG cells were exposed to ligands under the same conditions as in (Figure 2A) for 6h, and mRNA levels were assessed using RNA sequencing. The data contains n=4 biological repeats. The expression is plotted for BMP (A) and TGFβ (B) responsive genes. (C) Frequency distribution of the directional shift (*Log*_2_ *FC*_*Shift*_) from the additive model for the 61 BMP4-responsive (purple) and 783 TGFβ1-responsive (green) genes. The dashed lines represent the median shift for each group. The baseline (0) is represented by the gray vertical line. (D) EMT score as determined by PCA analysis of EMT- and MET-related genes (Table S2). The data contains n=4 biological repeats. (E) WT NMuMG cells were cultured for 36h under four different conditions: no ligands added (None), BMP4 at 100 ng/ml (BMP4), TGFβ1 at 1 ng/ml (TGFβ1), and a combination of BMP4 and TGFβ1 at these concentrations (TGFβ1+BMP4). The EMT state of the cells was determined by brightfield imaging and immunofluorescence staining of actin filaments with phalloidin (green) and E-cadherin levels (magenta). DAPI was used to visualize cell nuclei (blue). Scale bar, 20 μm. In all figures, *P ≤ 0.05, **P ≤ 0.01, ***P ≤ 0.001, ****P ≤ 0.0001, determined using a one-sided t-test in panels A,B and two-sided t-test in panel D. See also Figure S4.

To study the crosstalk on a genome-wide scale, we fitted the data to a generalized linear model (GLM) including an interaction term between the two ligands (see Methods). We defined a directional shift metric (*Log_2_FC_shift_*), quantifying the deviation from an additive model by comparing the observed response under combined stimulation to the response expected in the absence of an interaction term, such that positive values indicate synergistic enhancement and negative values signify antagonistic suppression. We applied this metric to genes responsive exclusively to either BMP4 or to TGFβ1. The distribution of these shifts revealed a global, asymmetric deviation from the additive model (Figure 3C, S4A). While genes responsive exclusively to BMP4 exhibited a robust and consistent negative shift, indicating widespread antagonism, TGFβ1-responsive genes showed a modest positive, though statistically significant, shift (Figure S4B,C). This smaller effect size in TGFβ1-responsive genes likely reflects that the mixed condition closely resembles the TGFβ1-induced transcriptional program, such that the addition of BMP4 produces only a modest change. In contrast, BMP4-responsive genes exhibit a stronger shift, as the mixed condition drives the system toward a distinct, TGFβ-like transcriptional program. The global relationship was further studied through Principal Component Analysis (PCA) of the entire responsive gene set, which shows that the measured mixed-signaling state deviates significantly from an additive model. Furthermore, the mixed response shows a TGFβ1-like PCA signature, further demonstrating that the pathway resolves the ambiguity by biasing towards the TGFβ1 transcriptional response (Figure S4D). Together, these analyses demonstrate that the asymmetric crosstalk operates at the genome-wide level, producing a biased, non-additive transcriptional response that resolves ambiguous signaling inputs towards TGFβ-like responses.

To assess the effect of early perception of signals on cellular decision, we analyzed EMT progression in NMuMG epithelial cells. From our RNA sequencing data, we curated a set of 92 EMT- and MET-related markers (Table S2)^44,49–51^. PCA performed specifically on these markers revealed distinct clustering based on the ligand environment (Figure S4E). While control and BMP4-treated cells cluster together in the epithelial regime, the addition of BMP4 to TGFβ1 drives the cells further along the first principal component compared to TGFβ1 alone, effectively enhancing the EMT signature. We used the first component from the PCA analysis of the curated set, which captures the maximal variation between signaling conditions, as an EMT score (Figure S4F,G). Classical mesenchymal markers such as *Snai1*, *Snai2*, *Vim*, *Fn1*, *Zeb1*, *Zeb2*, and *Serpine1*, contribute positively to the score, while epithelial markers like *Cdh1*, *Tjp1*, *Klf5*, *Cdn18*, and *Krt19* have negative contributions. We find that TGFβ1 induction significantly increases the EMT score (Figure 3D), while BMP4 alone has no effect. However, when supplied together with TGFβ1, BMP4 enhances the score beyond the levels induced by TGFβ1 alone.

To further validate the effect of the initial gene expression pattern, we analyzed the long-term phenotypic response in both 2D monolayer cultures and 3D organoid systems. We cultured NMuMG cells for 36 hours and determined EMT progression through staining of Phalloidin and E-cadherin (Figure 3E, S4H). Treating the cells with either BMP4 or a low level of TGFβ1 did not lead to a reduction in the levels of E-cadherin, a central epithelial marker, nor did it lead to the formation of stress fibers. However, cells treated with low amounts of TGFβ1 together with BMP4 showed a significant loss of E-cadherin, formation of stress fibers, and elongation of the cells and nuclei, all indicative of EMT transition. Using qPCR, we compared the response of *Serpine1*, a direct TGFβ target, and found that under mixed conditions, its levels are comparable to those induced by an EMT-inducing high-dose of TGFβ1 (4 ng/ml) (Figure S4I,J). This suggests that BMP4-mediated amplification of TGFβ signaling is sufficient to induce the EMT program. Finally, we stimulated 3D human breast organoid cultures with BMP and TGFβ. Under these conditions, co-stimulation with BMP4 and TGFβ1 enhances EMT-like features relative to TGFβ1 alone (Figure S4K,L). These findings indicate that the BMP-TGFβ asymmetric crosstalk potentiates EMT in both 2D and 3D models, such that BMP amplifies TGFβ-induced transcriptional and morphological hallmarks of EMT.

### BMP-TGFβ asymmetric crosstalk arises independently of feedback mechanisms

Next, we aimed to identify the molecular mechanism underlying the combinatorial crosstalk between the pathways. Feedback regulation is a common feature of many signaling networks, and BMP-TGFβ in particular, where ligand stimulation can induce the expression of transcription factors, signaling inhibitors, and other regulatory proteins that modulate pathway activity over time. To investigate whether feedback regulation underlies the asymmetric crosstalk, we first examined the temporal dynamics of the transcriptional response. Our fluorescent reporter measurements of BMP and TGFβ pathway activity were taken after 24 hours, by which time feedback effects may have begun to influence the response. We therefore performed a time-course analysis of the accumulation of fluorescence at multiple time points (Figure S5A). The fluorescent signal can be observed as early as 3 hours post-stimulation, maintaining a consistent crosstalk pattern across all timepoints throughout the 24-hour period. The early onset and stability of the asymmetric crosstalk suggest that it is not the result of transcriptional feedback.

To directly test the involvement of de novo protein synthesis, we tested the response of cells to ligand stimulation in the presence of Cycloheximide (CHX), a potent translation inhibitor. By preventing the production of new proteins, CHX effectively blocks transcriptional feedback loops, ensuring that any observed signaling alterations arise from primary pathway activation rather than secondary regulatory effects. When exposed to CHX, we find that the accumulation of both mCherry and mCitrine fluorescent reporters is abolished (Figure S5B), demonstrating the lack of newly synthesized proteins in our cells. Furthermore, measuring the mRNA levels of our fluorescent reporters reveals that the crosstalk pattern at the transcriptional level persists (Figure S5C). Consistent with our previous observations of the crosstalk, *mCherry* (BRE) transcript levels were depleted under BMP4 and TGFβ1 co-stimulation compared to stimulation with BMP4 alone. Similarly, *mCitrine* (TRE) transcripts were elevated relative to treatment with TGFβ1 alone. This asymmetry was observed both in the presence and absence of CHX, demonstrating that the asymmetric crosstalk between BMP and TGFβ signaling is established independently of transcriptional feedback or de novo protein production.

### SMADs phosphorylation levels directly reflect the ligand environment of the cells

To determine how the protein interaction network leads to asymmetric crosstalk, we analyzed the structure of the pathway (Figure 4A) and identified two levels within the pathway where the crosstalk could arise. Extracellularly, cross-interactions between ligands and receptors, as they combine to activate the SMAD mediator, could exert complex effects on the pathway response^37^. Alternatively, the crosstalk might arise intracellularly, downstream of the SMAD mediators, as they trimerize and drive the activation of target genes. To distinguish between these two levels, we directly examined the phosphorylation levels of the BMP-related SMAD1/5/8 and the TGFβ-related SMAD2/3 in response to ligand stimulation. For each pathway, we compared the measurement of the transcriptional response (see Figures 2D-F) to the phosphorylation levels of the SMAD second messengers (Figures 4B,C). When exposing cells to four concentrations of BMP4, we observed a dose-dependent increase in SMAD1/5/8 phosphorylation (Figure 4B). We next exposed cells to the same BMP4 concentrations but with the addition of TGFβ1 at a fixed concentration. Despite a substantial reduction in transcriptional activity under these conditions (Figure 2D), the phosphorylation levels of SMAD1/5/8 remained unchanged at both early (45 minutes) and later (6 hours) time points (Figure 4B). Similarly, we measured the levels of phosphorylated SMAD2/3 in cells exposed to TGFβ1 across concentrations, with or without BMP4 (Figure 4C). As before, while transcriptional activity levels increased under BMP4 co-stimulation (Figure 2E), phosphorylation levels of SMAD2/3 remained unaffected at both early and later time points. These findings demonstrate that the phosphorylation of the SMAD proteins faithfully reflects the concentration of their corresponding ligand, with no crosstalk between the pathways. Thus, we concluded that the crosstalk observed at the transcriptional level emerges downstream of SMAD phosphorylation.

**Figure 4.**
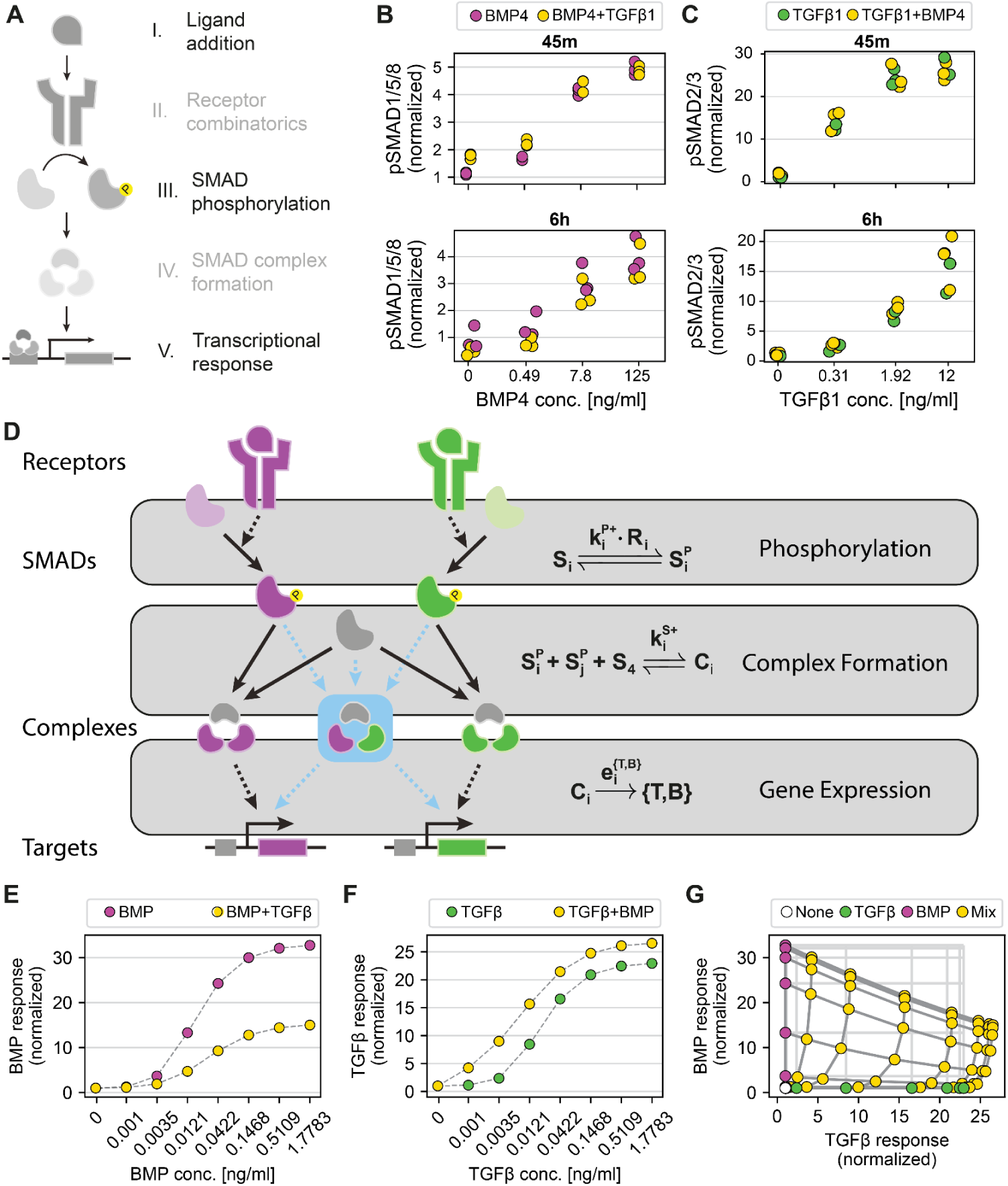
Asymmetric crosstalk resolves signaling ambiguity downstream of SMAD phosphorylation. (A) Schematic depiction of the different stages of signaling along the two pathways from ligand addition to transcriptional response. (B) Phosphorylated SMAD1/5/8 levels were determined by antibody staining after 45m and 6h of exposure to four concentrations of BMP4 across its dynamic range, with (yellow) and without (purple) addition of TGFβ1 at 1.92 ng/ml. Cf. Figure 2D. (C) Phosphorylated SMAD2/3 levels were determined by antibody staining after 45m and 6h of exposure to four concentrations of TGFβ1 across its dynamic range, with (yellow) and without (green) addition of BMP4 at 31.25 ng/ml. Cf. Figure 2E. (D) Schematic depiction of a mathematical model for the BMP and TGFβ pathways. The model comprises kinetic equations for SMAD phosphorylation, complex formation, and target gene expression. (E-G) Simulations for one parameter set showing asymmetric crosstalk (parameters are given in Table S3), showing the pathways’ response to concentration gradients of BMP (E) or TGFβ (F) ligands (purple or green, respectively), compared to the same gradients together with a constant ligand from the opposite pathway (yellow). Cf. Figures 2D-F. (G) Simulations of the full two-dimensional ligand gradient with BMP and TGFβ ligands. The BMP response and TGFβ response are plotted for all combinatorial conditions, including no ligand (white), BMP ligand only (purple), TGFβ only (green), and mixed conditions (yellow). The light gray grid indicates the expected response coordinates of a mixed sample, assuming no crosstalk effects. For all panels in this figure, the data show n=3 biological repeats. See also Figure S5.

### Promiscuous SMAD complex formation explains the observed asymmetric crosstalk

Our experimental data have revealed a biased asymmetric crosstalk between the BMP and TGFβ pathways, which emerges downstream of the SMAD phosphorylation. To uncover the possible mechanism underlying this crosstalk, we developed a mathematical model describing the known intracellular interactions in the BMP and TGFβ pathways (see Supplementary Information). The model describes ligand-dependent phosphorylation of SMADs, trimeric SMAD complex formation, and the downstream expression of target genes (Figures 4D and S5D). For simplicity, we consider all SMAD isoforms within each pathway to be equivalent and denote all variants as S_1_ for the BMP pathway (SMAD1, 5, and 8) and S_2_ for the TGFβ pathway (SMAD2 and 3). The overall SMAD phosphorylation rate is proportional to the number of receptor-ligand complexes (R_1_ for BMP and R_2_ for TGFβ) and the phosphorylation rate by an individual receptor, denoted by k^p^_1_ and k^P^_2_. Phosphorylated SMADS (S^p^_1_, S^p^_2_) can then form trimeric complexes, C_1_ and C_2_, which are composed of two S^P^_1_ or two S^P^_2_ molecules, respectively, and a single co-SMAD, S_4_. The binding affinities for the formation of C_1_ and C_2_ are denoted as k^C^_1_ and k^C^_2_. We further consider the trimeric complexes to induce the expression of a BMP target, B, with transcriptional efficacy of e^B^_1_, and the TGFβ target, T, with efficacy of e^T^_2_. This 11-parameter model captures the known pathways architecture and provides a quantitative framework to study the crosstalk computationally.

Using this model, we analyzed possible patterns of crosstalk across distinct parameter regimes. For each parameter set, we determined the crosstalk by simulating the pathway response for combinations of ligands, represented by the values of R_1_ and R_2_ (Figure S5E). Using dimensional analysis, we find that 5 parameters simply rescale the response and do not affect the crosstalk pattern (see Supplementary Information). Thus, to identify all obtainable crosstalk patterns, it is sufficient to keep these 5 parameters fixed and scan over the remaining 4 parameters (the levels of the S_1_, S_2_, and S_4_, and the ratio between the affinities of forming the two complexes). Intriguingly, we found that the model gives rise to only a mutual suppressive effect on both pathways across all parameters (red region) and does not recapitulate the experimental observations of an asymmetric crosstalk (yellow region) (Figure S5F). This purely antagonistic crosstalk stems from the competition between the two pathways over the shared SMAD4 during complex formation. While the competition for SMAD4 can explain the decreased BMP response, it cannot account for the increased response observed in the TGFβ pathway. Thus, in its current form, the model fails to reproduce the experimentally observed asymmetric crosstalk.

Attempting to capture the observed asymmetric behavior, we extended our model to allow combinatorial interactions between SMADs from the two pathways (Figure 4D, blue box). This extension enabled the formation of heteromeric SMAD complexes comprising both S_1_ and S_2_ (see Supplementary Information). Indeed, the existence of such complexes has been discussed previously^31,34,52^. The response structure in this extended model is governed by the four parameters of the canonical model, along with three additional ones, the affinity of forming the heteromeric complex (k^c^_mix_) and the two efficacy parameters for the heteromeric complex (e^B^_mix_, e^T^_mix_). Systematic simulations across these seven parameters reveal diverse crosstalk patterns (Figure S5G). In addition to mutual antagonistic crosstalk (red region), we identified a mutual synergistic regime (green region) and two asymmetric crosstalk regimes (yellow regions). We also note that the extended model enables more significant inhibitory effects, even within the mutual antagonistic regime. Importantly, one of the asymmetric crosstalk regimes reflects our experimental observations, where the addition of TGFβ decreases the BMP response across concentrations, while BMP addition results in an increased TGFβ response (Figures 4E,F). Furthermore, simulating varied concentrations of the two ligands reproduces the deformation of the BMP-TGFβ response grid observed experimentally (Figure 4G), reinforcing the model’s validity. We note that while the qualitative features of the asymmetric crosstalk are well captured, the quantitative response at low BMP or low TGFβ signals slightly deviates from the data. These deviations may result from our assumption of a linear transcriptional response. However, cooperative effects and others generally lead to nonlinear responses, which can recapitulate these characteristics in the data (see Supplementary Information and Figure S6A). Thus, we find that a mixed SMAD complex can provide a mechanism for reproducing the experimentally observed asymmetric crosstalk.

### Systematic parameter analysis and experimental validation for heterocomplex-mediated crosstalk

The parameter set used to simulate the asymmetric crosstalk in Figure 4E-G is representative, but not unique. To better understand the conditions resulting in asymmetric behavior and test the robustness of our model, we performed a comprehensive scan of the parameter space and systematically plotted both one-dimensional and two-dimensional distributions of these parameter sets (Figure S6B,C). Overall, we find that asymmetric crosstalk emerges broadly across diverse combinations of SMAD levels and complex affinities, rather than requiring a narrowly tuned regime. Furthermore, the asymmetric behavior arises from combinations of parameters, which can independently span wide values, rather than from a single specific value, underscoring the generality of the mechanism. By analyzing the resulting distributions (Figure S6C,D), we were able to identify relevant effects for specific parameters, some of which result in testable predictions for our model.

The basic premise of the model is the existence of heteromeric complexes, and the corresponding parameter is the affinity for the formation of these complexes, k^C^_mix_. As can be expected, we find that the higher k^C^_mix_ is, the more likely it is for an asymmetric response to arise. To explicitly study the role of this parameter in the model, we simulated the crosstalk across increasing affinity values (Figure 5A). Simulations with low heteromeric complex affinity showed a large overlap region where both pathways are active. Increasing the heteromeric complex affinity resulted in separated regions where only one pathway is active, as was seen experimentally. The existence of such complexes and their inhibitory effect has been discussed previously^31^. Our model suggests a new mechanism that utilizes these heteromeric SMAD complexes to asymmetrically enhance TGFβ signaling while suppressing BMP signaling, thereby reducing the ambiguity at the level of the transcriptional response.

**Figure 5.**
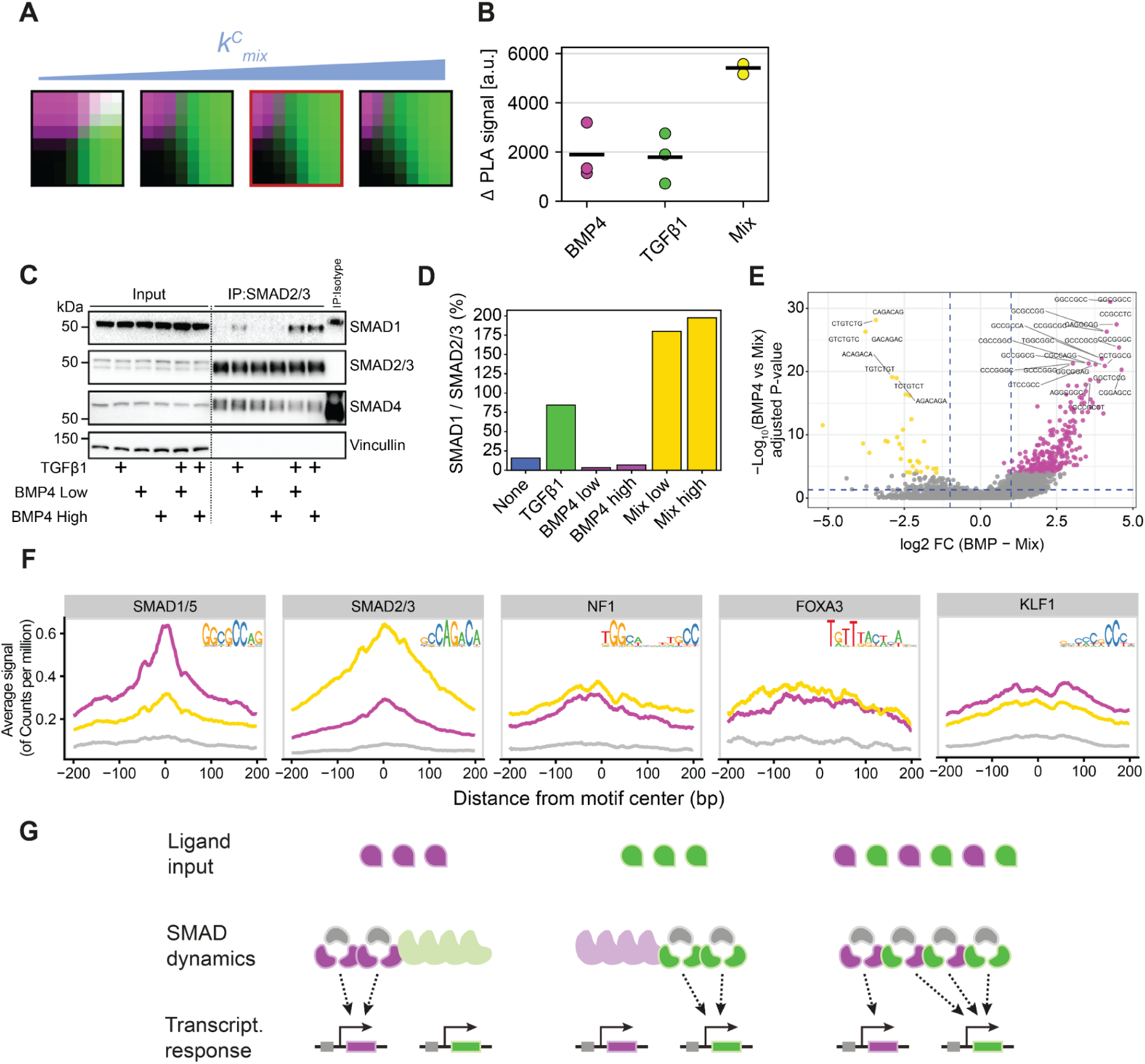
Asymmetric BMP–TGFβ crosstalk via heteromeric SMAD complexes shapes cell fate decisions. (A) Depiction of cell response space of two-dimensional concentration gradient assays with varying k^C^_mix_ and resulting changes in pathway-specific cell fate regions. The parameter set outlined in red represents the parameters used in Figure 4E-G. (B) NMuMG cells were exposed to four different conditions: no ligand, BMP4 at 31.25 ng/ml (BMP4), TGFβ1 at 1.92 ng/ml (TGFβ1), or a mix of both (Mix). Proximity ligation assay (PLA) was used to quantify the interaction of phosphorylated SMAD1/5/8 with phosphorylated SMAD2/3 under each condition. Data shown is the difference between the PLA signal and the no ligand control. (C) NMuMG cells were exposed to six different conditions: no ligand, TGFβ1 at 1.92 ng/ml, BMP4 at 7.8 ng/m (BMP4 low) and 31.25 ng/ml (BMP4 high), TGFβ1 at 1.92 ng/ml with BMP4 at 7.8 ng/ml (Mix low), and TGFβ1 at 1.92 ng/ml with BMP4 at 31.25 ng/ml (Mix high). Western blot analysis of SMAD1, SMAD2/3, and SMAD4 following immunoprecipitation of SMAD2/3. Whole lysates (input) are shown as controls. Vinculin serves as a loading control. (D) Densitometric analysis of Co-IP results from (C), showing the percentage of SMAD1 co-precipitating with SMAD2/3 under the indicated ligand conditions. (E-F) NMuMG cells expressing SMAD1 fused to MNase were treated with four different conditions: no ligand, TGFβ1 at 1.92 ng/ml, BMP4 at 48 ng/ml (BMP4), and a mix of both (Mix). (E) Volcano plot showing differential 7-mer enrichment in genomic regions comparing BMP4 and Mix conditions. Sequences significantly enriched in the BMP4 condition are highlighted in purple, while those enriched in the Mix condition are in yellow. (F) Read density profiles centered on specific transcription factor motifs (SMAD1/5, SMAD2/3, NF1, FOXA3, and KLF1). The profiles show binding intensity for BMP4 (purple), Mix (yellow), and non-treated (gray) samples. Relevant sequence logos are shown for each motif. (G) Schematic depiction of the crosstalk mechanism in the model. (left) When only BMP ligand is provided, three units of BMP ligand phosphorylate four units of BMP-related SMADs, which leads to the creation of two homogeneous BMP-SMAD complexes. The complexes then induce a transcriptional response of the BMP target genes. (middle) When only TGFβ is provided, three units of TGFβ ligand phosphorylate four units of TGFβ-related SMADs, which leads to the creation of two homogeneous TGFβ-SMAD complexes. The complexes then induce a transcriptional response of the TGFβ target genes. (right) When both BMP and TGFβ ligands are provided simultaneously, three units of BMP and three units of TGFβ result in four units of BMP-related SMADs and four units of TGFβ-related SMADs (no crosstalk). When both pathways are activated, some heterogeneous complexes are formed. When the TGFβ-related SMADs show higher affinity to the DNA targets, these heterogeneous complexes will preferentially bind to the TGFβ targets, increasing their expression. In addition, fewer homogeneous BMP-SMAD complexes are formed, reducing the expression of the BMP targets. Data shown contains n=3 biological repeats. See also Figure S6.

To experimentally test potential SMADs heteromerization, we utilized the Proximity Ligation Assay (PLA)^53,54^. This technique enables the detection of stable interactions between two proteins of interest, yielding a fluorescent signal measured using flow cytometry^55,56^. Our model predicts the existence of mixed TGFβ-BMP SMAD complexes in cells exposed to both ligands. We thus cultured cells with no added ligands, with TGFβ1 alone, with BMP4 alone, and with a mix of the two ligands. Following 24 hours of culture, we performed PLA using antibodies against SMAD1/5/8, the BMP-related SMADs, and against SMAD2/3, the TGFβ-related SMADs, and examined potential binding between the two groups. Consistent with our model predictions, we observed a strong signal only in cells cultured with both BMP4 and TGFβ1 (Figure 5B). To further validate the formation of heteromeric complexes, we performed co-immunoprecipitation (Co-IP) using an antibody against SMAD2/3 followed by immunoblotting for SMAD1. While SMAD1 was largely absent from SMAD2/3 immunoprecipitated samples in untreated cells or cells treated with either ligand alone, a robust interaction was observed under combined BMP4 and TGFβ1 stimulation (Figure 5C,D), consistent with the PLA results and our model predictions. We note that while we find an increased interaction between SMAD1/5/8 and SMAD2/3, it could also occur through higher-order structures. Additionally, as the methods are qualitative, it remains unclear what fraction of the activated SMAD trimers in co-treated cells are heterotrimers. However, these findings show that simultaneous activation of the two signaling pathways results in increased interactions between SMAD1/5/8 and SMAD2/3.

The cofactor SMAD4 is a central component for SMAD complex formation. To determine whether SMAD4 abundance modulates the crosstalk, we analyzed the model’s behavior across a range of levels for the corresponding parameter, S_4_. We found that the asymmetric regime persists when S_4_ is present at levels comparable to or exceeding S_2_ (Figure S6C,D). Thus, the model predicts that SMAD4 is not a limiting factor and the inhibition arises from the formation of mixed complexes rather than competition for SMAD4 molecules. Indeed, it has been shown that in biological contexts, SMAD4 is commonly abundant, in agreement with the model^31,57,58^. To further validate its abundance in our cells, we performed siRNA-mediated knockdown of SMAD4. We found that a reduction of over 50% in SMAD4 levels did not affect the cellular response to either BMP or TGFβ, supporting the model’s prediction of abundant SMAD4 and aligning with previous observations that partial depletion of SMAD4 leaves many TGFβ responses intact^59^. Accordingly, under combined BMP4 and TGFβ1 stimulation, the response of TGFβ remained enhanced while the BMP response was suppressed (Figure S6E). These findings showcase the robustness of the asymmetric crosstalk and support the model’s prediction by demonstrating the abundance of SMAD4.

Our model also shows that efficacy parameters of the SMAD complexes critically affect the crosstalk. Parameter analysis reveals that the key parameters are the fold change difference in the efficacy between a heteromeric and a homomeric complex, e^B^_mix_/e^B^_1_ and e^T^_mix_/e^T^_2_. Analyzing the simulation results, we find that a TGFβ-biased asymmetric crosstalk is more likely to arise when the BMP efficacy fold-change is lower than the TGFβ efficacy fold-change. Intuitively, this means that the mixed complex behaves more like a SMAD2/3 complex than like a SMAD1/5/8 complex. This suggests that by forming the mixed complexes, BMP-associated SMAD proteins are mostly being used to activate TGFβ targets, enhancing the TGFβ pathway activity.

Several biological processes contribute to the efficacy of a transcription factor, including DNA binding affinity, transcription initiation rate, and the number of target sites. The efficacy parameter combines all these processes into an effective parameter representing the overall expression rate of a single BMP or TGFβ target regulated by a specific complex. Accordingly, one mechanism to increase efficacy value is through an increased affinity between the complex and the target sites. Affinity dependence would result in a gene-specific crosstalk pattern. In our case, we find that the asymmetric crosstalk occurs globally across different target genes (see Figure 3C), suggesting that the binding affinity has a minor effect.

Alternatively, differences in the number of target sites for SMAD1/5/8 vs SMAD2/3 could give rise to a global effect on the efficacies. Complexes with more target sites spend, on average, less time bound to each single target, resulting in lower average expression of each individual gene and a lower efficacy. Assuming that the mixed complexes bind both targets of SMAD1/5/8 and of SMAD2/3, heteromeric complexes will acquire many more targets compared to a SMAD1/5/8 homomeric complex. This will globally reduce the relative BMP efficacy more than it would affect the relative TGFβ efficacy, in agreement with the model prediction. Our RNA sequencing dataset shows a higher number of TGFβ1-responsive genes (783) compared to BMP4-responsive genes (61). Previous ATAC-seq studies ^60,61^ show that while there are similar numbers of overall BMP and TGFβ sites in NMuMG cells, the TGFβ sites are in open chromatin regions, while a significant amount of the BMP sites require chromatin remodeling. This could provide an explanation for some of the efficacy differences, together with other biochemical parameter differences.

Finally, our model predicts that mixed complexes induce TGFβ targets by altering the genomic targeting of BMP-associated SMAD proteins. This results in the redistribution of BMP-associated SMAD proteins towards TGFβ-binding motifs. To test this prediction, we performed ChEC-Seq ^62,63^ and determined the genomic localization of SMAD1 under different conditions (Figure S6F-H). At the global level, we quantified differential SMAD1 occupancy across the genome and classified peaks based on their relative signal between BMP4 and mixed (BMP4 and TGFβ1) conditions (Figure S6I). We identified 402 peaks with enhanced occupancy under BMP4 stimulation and 637 peaks with enhanced binding under mixed stimulation (Figure S6I,J). We next performed differential k-mer enrichment analysis between these two peak sets to identify sequence features (7-mers) associated with condition-specific binding (Figure 5E). This analysis revealed distinct enrichment patterns. Sequence features enriched under BMP4 stimulation were associated with canonical SMAD1/5 motifs (GGCGCC-like), whereas features enriched under mixed stimulation were associated with canonical SMAD2/3-associated motifs (CAGA-like). To further characterize the redistribution of SMAD1 binding, we analyzed the average ChEC-seq signal around motif centers, found using de novo motif analysis (Figure 5F). While SMAD1/5 and SMAD2/3 motifs showed distinct, condition-specific enrichment, other motifs (NF1, FOXA3, and KLF1) exhibited non-specific binding across conditions. Specifically, SMAD1 occupancy at SMAD1/5 motifs was high under BMP4 stimulation and drastically reduced under mixed conditions to levels similar to non-specific motifs, whereas the signal at SMAD2/3 motifs was specifically increased under mixed stimulation. Together, these results show that SMAD1 binding is redistributed under mixed signaling conditions toward regions associated with TGFβ-responsive transcription, providing genome-wide support for the model prediction that heteromeric SMAD complexes redirect BMP-associated SMADs to TGFβ target genes.

To assess the functional consequences of SMAD1 redistribution, we integrated the ChEC-seq binding data with our RNA-seq gene expression data. We identified ChEC-seq peaks located near gene promoters and classified them based on their condition-specific enrichment, distinguishing peaks enhanced under BMP4 from those enhanced under mixed stimulation. Projecting these peak categories onto a differential expression volcano plot of the RNA-seq data (Figure S6K) revealed a strong correspondence between SMAD1 genomic occupancy and transcriptional output. Genes that were significantly upregulated under BMP4 stimulation were associated with BMP4-enhanced peaks near their promoters, whereas genes upregulated under mixed conditions were associated with peaks enhanced under mixed stimulation.

Overall, these results validate our model’s predictions and identify mixed SMAD complexes as key mediators of asymmetric signal integration. These complexes act to suppress BMP output and amplify TGFβ responses, thereby resolving pathway ambiguity through biased transcriptional efficacy and genomic targeting. Our model offers an intuitive understanding of the molecular mechanism underlying this asymmetric crosstalk (Figure 5G). The asymmetric crosstalk arises in a parameter regime where the formation affinity of the heteromeric complex (k^C^_mix_) is high, and its expression rates are skewed towards one of the pathways (e^B^_mix_ / e^B^_1_<< e^T^_mix_ / e^T^_2_). In this scenario, when cells are exposed to BMP ligands, only SMAD1/5/8 are phosphorylated and bind to form complexes, leading to the activation of BMP target genes. Similarly, TGFβ ligands activate the SMAD2/3 proteins and, consequently, the TGFβ target genes. However, the exposure of cells to environments containing ligands of the two pathways results in the formation of SMAD heteromers. Such heteromers induce the expression of TGFβ targets at higher rates. Thus, the TGFβ pathway utilizes the BMP-associated SMADs and reroutes them to activate TGFβ targets, skewing the response in favor of TGFβ.

## Discussion

Signaling pathways that regulate cell fate often operate in opposition to one another, posing a challenge for cells that must resolve contradictory cues into coherent responses. The BMP and TGFβ pathways often exhibit conflicting effects on cellular behavior and cell fate decisions^17–25^. Nevertheless, in various developmental and pathological contexts, such as gastrulation, germ-line formation, and cancer progression, cells frequently encounter ligands that induce the simultaneous activation of both pathways^4–6^. Understanding how cells resolve such combinatorial signals is key to decoding cell fate decisions and gaining predictive control over developmental systems.

In this study, we systematically examined the cellular response to simultaneous exposure to BMP and TGFβ signals. While prior studies have demonstrated mutual inhibition between these two pathways^22,31,46,64^, our findings reveal a more nuanced form of crosstalk. While the BMP response is inhibited, the TGFβ signaling is enhanced, leading to transcriptional activity dominated by the TGFβ program. This asymmetric crosstalk is evident across multiple cell types, ligand variants, and concentrations, and allows cells to resolve the ambiguity of simultaneous BMP and TGFβ signaling.

The finding that TGFβ and BMP ligands affect both the amplitude and EC_50_ of the opposing pathway highlights a complex mode of crosstalk. Rather than a simple inhibition or enhancement, the crosstalk reshapes the dose-response properties of each pathway. TGFβ reduces the BMP pathway’s maximal output and increases its EC_50_, thus decreasing both its amplitude and its sensitivity. Conversely, BMP increases the amplitude of the TGFβ response and slightly lowers its EC_50_, enhancing both its strength and sensitivity. These effects may explain the behavior of BMP10, which activates both pathways and shows a reduced saturation level of the BRE and an increased saturation level of the TRE. Nonetheless, this work primarily focused on ligands that activate only one of the pathways, in order to gain a clearer mechanistic understanding of how the TGFβ and BMP pathways interact.

Interestingly, we detected differences in the crosstalk intensity among various ligand pairs, even though all ligands were applied at similar effective concentrations. This variability may stem from differences in SMAD protein usage (e.g., SMAD1 vs. SMAD5 or SMAD2 vs. SMAD3), receptor binding affinities, or cofactor recruitment. Future studies could explore how different ligand-receptor-SMAD combinations modulate the formation or activity of downstream complexes.

To uncover the mechanism behind the asymmetric crosstalk, we utilized a combination of mathematical models and experimental measurements. Our mathematical model predicted that the formation of heteromeric SMAD complexes, containing components from both the BMP and TGFβ pathways, could explain the observed transcriptional bias. These complexes redirect BMP-associated SMADs toward TGFβ target genes, suppressing BMP output while enhancing TGFβ responses. The formation of such complexes was previously shown in response to BMP7 and TGFβ1 to have an inhibitory effect on BMP signaling^31^. Our work extends this phenomenon, demonstrating that the heteromeric complexes exhibit inhibitory effects across various BMP and TGFβ ligands. Remarkably, we find that mixed SMAD complexes can also act as functional enhancers of TGFβ activity. Thus, our results propose that the formation of these mixed complexes redirects the SMAD components of the BMP pathway to activate TGFβ targets, thereby favoring TGFβ-like responses.

Indeed, we validated these predictions experimentally using complementary biochemical and genomic approaches. Using PLA and co-immunoprecipitation, we show that SMAD1 and SMAD2/3 form mixed complexes specifically under mixed ligand conditions. In parallel, ChEC-seq reveals a redistribution of SMAD1 binding from canonical BMP-associated motifs toward TGFβ-associated motifs under co-stimulation, further supporting the notion that heteromeric complexes redirect transcriptional specificity. While our data support the model predictions, the precise composition of the complexes occupying these sites remains to be fully resolved. In particular, the baseline binding of SMAD1 at TGFβ-associated motifs under BMP4 stimulation raises the possibility that SMAD1 homomeric complexes may also bind these sites, potentially with low affinity or in a context-dependent manner. Under mixed stimulation, increased accessibility of TGFβ-responsive elements or recruitment of additional cofactors could further enhance this binding. The current data do not exclude a contribution from SMAD1 homomeric binding at TGFβ-associated sites. Importantly, both scenarios could combine to shape the crosstalk structure. Future work aimed at directly resolving the composition of SMAD complexes at specific genomic loci will be required to distinguish between these possibilities.

TGFβ signaling is often associated with the acquisition of cellular plasticity^59,65^, while BMP typically directs cellular differentiation^5,19,23^. Our findings suggest that under complex signaling environments, the architecture of these pathways resolves ambiguity in a manner that promotes plasticity by favoring TGFβ-like responses. This adaptation may benefit cells in dynamic environments by providing them with the ability to adapt, regenerate, and respond to environmental changes, a crucial feature in various biological processes and disease contexts^66,67^.

The observed crosstalk exhibits several distinctive properties. First, the asymmetry manifests exclusively in combinatorial contexts. When cells are exposed to each ligand individually, we observe distinctly orthogonal activation responses. Only when both pathways are simultaneously active do we observe their mutual influence. Second, we find that the nature of the response to specific signals is significantly influenced by the overall extracellular signaling milieu, as indicated by recent studies^68^. When the TGFβ pathway is active, the cellular state transforms the role of a BMP signal from activating canonical BMP targets to activating canonical TGFβ targets. A TGFβ-dependent role of BMP was previously shown to occur in the induction of regulatory T cells differentiation, where BMP by itself does not induce regulatory T cells but synergizes with TGFβ to induce their differentiation^69^. In our system, we find that when administered together with TGFβ1, BMP4 further drives the EMT progression of cells at both the transcriptional and phenotypic levels. These findings demonstrate that the asymmetric crosstalk between BMP and TGFβ signaling is not only embedded at the molecular level but also exerts functional consequences, markedly enhancing EMT progression when both pathways are co-activated. It is imperative to further investigate how this biased encoding of mixed signals governs cellular decisions, such as EMT, and to what extent this signal perception determines cell fate. Furthermore, such context dependence could potentially elucidate previously observed paradoxical effects of BMPs in processes like EMT and cancer progression, where BMP ligands can either inhibit or promote plasticity depending on the signaling environment^5,70^.

While our findings focus on the BMP-TGFβ pathways, they may reflect a broader principle in signaling networks. Recent studies have shown that complex, multilayered, and promiscuous interacting networks in receptor-ligand binding can perform functional computations in cells^35–37^. Our findings extend this concept to the promiscuous dimerization of pathway mediators, showing that heteromeric complex formation among SMADs adds computational capacity to the pathway. Importantly, this feature is not exclusive to the BMP and TGFβ pathways. For instance, the JAK/STAT pathway exhibits similar activation-dependent dimerization of the STAT second messengers^71–73^. In signaling pathways, it is often suggested that numerous ligands are encoded by only a few second messengers, implying that much information may be lost. However, our results propose that heteromeric complexes could introduce the necessary complexity required to resolve signal combinations and encode context-sensitive decisions. Understanding the generality of this principle could open up new avenues for addressing the general question of redundancy in signaling pathways.

## Methods

### Tissue culture and cell lines

NMuMG (NAMRU Mouse Mammary Gland cells, female) were purchased from ATCC (CRL-1636). ATDC5 cells were a generous gift from Prof. Michael Elowitz (Caltech). All cells were cultured in a humidity-controlled chamber at 37°C with 5% CO2. Cells were cultured in DMEM supplemented with 10% FBS, 1 mM sodium pyruvate, 100 unit/ml penicillin, 100 µg/ml streptomycin, 2 mM L-glutamine, and 1x MEM nonessential amino acids.

### Dual-reporter cell line construction

Construction of the reporter cell lines was carried out via simultaneous random integration of plasmids harboring either the BMP response element (BRE)^39^ or the TGFβ response element (TRE)^39,41^ in the enhancer region of a minimal CMV driving the expression of an H2B-mCherry or an H2B-Citrine protein fusion, respectively. NMuMG and ATDC5 cells were transfected using Lipofectamine LTX and PLUS reagent. After transfection, cells were selected with 200 μg/ml Hygromycin. All experiments were performed with clonal populations generated via limiting dilutions.

### Microscopy

#### Dual reporter cells imaging

NMuMG dual reporter cells were plated at 40% confluency in 96-well glass-bottom optical plates (CellVis) and cultured under standard conditions in DMEM without Phenol red, containing all ingredients as specified above for 24 hours. Media was then replaced with fresh media containing ligands at the required concentrations. 24 hours following ligand addition, cells were imaged using the CellDiscoverer 7 (Zeiss).

#### EMT state determination in NMuMG cells

NMuMG cells were plated at 30% confluency in 96-well glass-bottom optical plates, and cultured under standard conditions in a similar Phenol red-free DMEM for 24 hours. Media was replaced with fresh media containing ligands at the required concentrations. 36 hours following media replacement, cells were then imaged using the CellDiscoverer 7 microscope (Zeiss).

#### EMT state determination in Human tumor breast organoids

Patient-derived tumor breast organoids were overlaid with organoid culture medium containing Advanced DMEM DMEM/F12 (Thermofisher; 12634010), supplemented with 10mM HEPES (GIBCOTM; 15630-080), 2mM GlutaMAX (GIBCOTM; 35050-061), x1 Penicillin streptomycin (Biowest; L0022-100), 10% R-spondin-1-conditioned medium (RCM) produced from HEK293 HA–Rspo1–Fc cells (Cultrex® HA–R-spondin-1–Fc 293T cells; 3710-001-01), 10% Noggin-conditioned medium (NCM) produced from HEK293 cells stably transfected with pcDNA3-mouse NEO insert (to confer neomycin resistance; cells for NCM production were kindly provided by the Hubrecht Institute, Utrecht, The Netherlands), x1 B27 (ThermoFisher; 17504044)), 100 ng/ml A83-01 (Tocris Bioscience; 2939), 50 ng/ml EGF (PeproTech; AF-100-15). When organoids were first established or expanded, 10 mM Y-27632 (ROCK inhibitor; Sigma Aldrich; Y0503) was added to the medium. Medium was changed every 4 days, and organoids were passaged every few weeks using mechanical shearing with TrypLE Express (Invitrogen; 12605036). Whole organoids were suspended in BME, plated into 18-well μ-Slides (ibidi; 81816), and overlaid with the appropriate growth medium for the duration of the experiment (5–8 days). Control organoids were cultured in standard organoid medium. TGFβ1-treated organoids were cultured in EMT-optimized medium, consisting of organoid medium without A-83-01, as previously described ^74^, supplemented with TGFβ1. BMP4 only organoids were cultured in standard organoid medium supplemented with BMP4. Organoids in the TGFβ1+BMP4 group were cultured in an EMT-optimized medium supplemented with TGFβ1 and BMP4. Confocal and wide field fluorescent imaging was performed with a TCS SP8 Leica Confocal microscope, equipped with a Leica DFC9000 GT camera (Leica Microsystems, Nussloch, Germany). Image processing (i.e., Crope, pseudo-coloring, merge channels) was performed with LAS X software (3.7.2.22383).

### Stimulation of cells and flow cytometry

NMuMG dual reporter cells were plated at 30% confluency in 96-well plates and cultured under standard conditions (specified above) for 24 hours. Media was then replaced with fresh media containing ligands at the required concentrations. 24 hours following ligand addition, cells were prepared for flow cytometry as follows: cells were washed with PBS and trypsinized for 5 min at 37°C. Digestion was quenched by re-suspending the cells in HBSS with 2.5mg/ml Bovine Serum Albumin (BSA). Cells were then filtered with a 20µm mesh (Merck NANMN2010) and analyzed by flow cytometry (NovoCyte Quanteon, Agilent). All recombinant ligands were acquired from R&D Systems (Table S4).

### Immunofluorescence staining

#### Assessment of SMAD phosphorylation levels

NMuMG cells were plated at 50% confluency under standard conditions in 24-well plates. Following 24 hours of culture, cells were added with the indicated ligands and incubated at 37°C for 45 min. Cells were washed with PBS, trypsinized for 5 min at 37°C, and re-suspended in PBS. Cells were further fixated in 4% formaldehyde for 15 min at room temperature (RT). Following fixation, cells were washed with PBS and permeabilized with Triton X-100 (0.1%) for 15 min at RT. Cells were washed with antibody dilution buffer (ADB, PBS + 0.5% BSA) and re-suspended in ADB containing 1:40 dilution of the antibody against the phosphorylated form of SMAD2/3 (BD, clone O72-670) and 1:200 dilution of the primary antibody against the phosphorylated form of SMAD1/5/8 (Cell Signaling Technologies, clone D5B10). Cells were stained for 1h at RT. Following incubation, cells were washed with ADB and re-suspended with ADB containing a 1:2000 dilution of a secondary antibody labeled with Alexa Fluor 405 (Abcam, ab175660), and incubated for 30 min at RT. Finally, cells were washed with ADB, re-suspended in HBSS + 2.5mg/ml BSA, filtered with a 20µm mesh, and analyzed by flow cytometry (NovoCyte Quanteon, Agilent).

#### Assessment of EMT markers in NMuMG cells

Cells were washed with PBS, fixed with 4% Formaldehyde for 20 minutes at RT, and permeabilized with 0.1% Triton X-100 for 15 minutes. Following permeabilization, cells were blocked with 5% BSA in PBS for 20 minutes and incubated with anti-E-cadherin antibody (24E10 Rabbit mAb Alexa Fluor 488 Conjugate, Cell Signaling #3199 1:300 in ADB) for 1 hour at RT. After washing, cells were incubated with Phalloidin (iFluor 647-conjugated, Abcam ab176759, 1:1,000 in PBS) for 30 minutes at RT. Nuclei were stained with DAPI (20µg/ml) for 5 minutes.

#### Assessment of EMT markers in Human breast organoids

Organoids were fixed in 4% Paraformaldehyde (PFA) for 30 minutes, permeabilized with 0.3% Triton X100 in PBS for 30 min, then blocked with 3% BSA in PBST (0.01% Triton X-100 in PBS) for 1 hour. The organoids were incubated with anti-E-Cadherin (Rabbit mAb, Cell Signaling; #3195, 1:200 in 3% BSA in PBST), for 2 hours at RT, then for 1 hour at RT with a secondary antibody (goat anti Rabbit, Alexa Fluor 647 conjugated, Abcam; ab150083, 1:500 in 3% BSA in PBST) together with Phalloidin-iFluor 488 Reagent (Abcam; ab176753, 1:1000). Incubations were followed by three washes of 5 min with PBST. DNA staining was performed with DAPI (Invitrogen, D1306) for 10 min, and then organoids were covered with PBS.

### Proximity Ligation Assay (PLA)

Heteromerization between BMP- and TGFβ-related SMADs was examined using Duolink proximity ligation assay (PLA) according to the manufacturer’s instructions (Duolink® flowPLA Detection Kit - Green; Sigma-Aldrich; cat#: DUO94102-1KT). NMuMG cells were plated at 65% confluency in 24-well plates under standard conditions. Following 24 hours of culture, cells were supplemented with the indicated ligands for an additional 45 min. Cells were harvested using trypsin and supplemented with PBS. Cells were further fixated in 4% formaldehyde for 15 min at room temperature (RT). Following fixation, cells were washed with PBS and permeabilized with Triton X-100 for 15 min at RT. Cells were further washed with antibody dilution buffer (ADB: PBS + 0.5% BSA) and the supernatant was removed. Cells pellets were added with a blocking solution (Sigma-Aldrich) and incubated at 37°C for 1h. Cells were washed with ADB and stained with antibodies diluted in ADB as follows: 1:40 dilution of the antibody against the phosphorylated form of SMAD2/3 (BD, clone O72-670) and 1:200 dilution of the antibody against the phosphorylated form of SMAD1/5/8 (Cell Signaling Technologies, clone D5B10). Cells were stained for 1h at RT. Following incubation, cells were washed with ADB and then washed again with Duolink wash buffer. Cells were added with PLA probes (PLUS and MINUS), incubated at 37°C for 1h, and washed with Duolink wash buffer. Cells were then ligated for 30 min at 37°C, washed, amplified with Polymerase for 100 min at 37°C, and added with detection solution for 30 min at 37°C. Following detection, cells were washed and analyzed by flow cytometry (NovoCyte Quanteon, Agilent).

### RNA Preparation, Library Construction, and RNA Sequencing

Library preparation protocol was bulk-adapted from MARS-seq^75,76^. Briefly, cells were plated at 50% confluency in 24-well plates. Following 24 hours, cells were added with the indicated ligands and cultured for an additional 6 hours. Cells were lysed in a lysis/binding buffer (Life Technologies), and polyadenylated RNA was purified using Dynabeads oligo dT (Life Technologies). RNA from each sample was barcoded, and samples with similar overall RNA content were pooled together, up to eight samples in a pool. Pooled samples underwent second-strand synthesis and were linearly amplified by T7 in vitro transcription. Amplified RNA was fragmented and converted into a sequencing-ready library by tagging the samples with Illumina sequences during ligation, RT, and PCR. Libraries were quantified by Qubit and TapeStation, and sequenced using the Illumina NextSeq 500.

### Cycloheximide treatment

To inhibit protein translation, NMuMG dual reporter cells were treated with Cycloheximide (CHX, Sigma-Aldrich #C7698-1G) at a final concentration of 1 mg/ml. CHX was added simultaneously with ligand stimulation. Cycloheximide was maintained in the media during the entire ligand exposure period. Cells were then harvested after 6 hours for RNA extraction and qPCR analysis. Similarly, NMuMG dual reporter cells were incubated with CHX and ligands for 6 hours followed by analysis in flow cytometry, as described above in the methods.

### Quantitative PCR

Total RNA was extracted from cell lysate using the RNeasy mini kit (Qiagen), and cDNA was generated from 1μg of RNA using the qScript cDNA synthesis kit (Quanta) according to the manufacturer’s instructions. Primers and probes for specific genes (Table S5) were purchased from Sigma-Aldrich. Reactions were performed using 1:25 dilution of the cDNA synthesis product with Fast SYBR Green Master Mix (ThermoFisher Scientific). Cycling was carried out on a QuantStudio 5 thermocycler using an initial denaturing incubation of 95° for 20 seconds, followed by 40 cycles of (95° for 1 second, followed by 60° for 20 seconds). Each condition was assessed with three biological repeats, and each reaction was run at least in triplicate. Sdha was used to normalize gene expression between samples.

### siRNA transfection

NMuMG dual reporter cells were transfected with Stealth RNAi siRNA against mouse Smad4 (ThermoFisher Scientific, #1320003) delivered using Lipofectamine RNAiMAX Transfection Reagent (ThermoFisher Scientific, #13778075), according to the manufacturer’s instructions. In short, cells were plated at 40% confluency in 24-well plates for 24h and then added with a combination of two siRNA against Smad4 (MSS206436 and MSS206437, 30nM each) for an additional 24h. Following 24h of transfection, cells were added with ligands as indicated for an additional 24h, for a total transfection time of 48h. The effect of SMAD4 knockdown on BMP and TGFβ pathway activation was measured using flow cytometry. To measure the extent of silencing, total RNA was extracted after 48h of transfection, and mRNA levels of SMAD4 were determined by quantitative PCR.

### ChEC-seq

NMuMG cells were transfected with Smad1-MNase using Dharmafect kb reagent. After transfection, cells were selected with 200 ug/ml Hygromycin for 10 days. A polyclonal population was used for this experiment. Cells were plated at 80% confluency in a 6-well plate, with 3 repeats per condition. 16 hours after plating, ligands were added to the cells for 30 minutes. Cells were then washed 3 times with buffer A (20mM Hepes Buffer (Biological Industries #03-025-1B), 110mM Potassium acetate, 5mM Sodium acetate, 0.2 mM spermine, 0.5 mM spermidine, 1× cOmplete EDTA-free protease inhibitors [Roche, one tablet per 50 ml buffer], 1 mM PMSF, 1.5mM EGTA). Cells were permeabilized using 250 µL of buffer A without EGTA and 0.05% digitonin for 5 min. Next, 50 µL CaCl2 was added to a final concentration of 2mM to activate the Mnase and incubated for 2 min. The CaCl2 treatment was stopped by adding 100 µL of stop buffer (800mM NaCl, 40mM EDTA, 8mM EGTA, and 40% SDS) to the cell suspension. After this, the cells were treated with Proteinase K (0.05 mg/ml final concentration) at 55°C for 1 hr. An equal volume of Phenol-Chloroform pH = 8 (Sigma-Aldrich) was added, vigorously vortexed, and centrifuged at 17,000g for 15 min to extract DNA. The DNA was precipitated after the extraction with 2.5 volumes of cold 96% EtOH, 45 mg Glycoblue, and 20 mM sodium acetate at –80°C for>1 hr. Next, the tubes were centrifuged (17,000 g, 4°C for 10 min), the supernatant was removed, and the DNA pellet was washed with 70% EtOH. The DNA pellets were dried and resuspended in 30 µL RNase A solution (0.33 mg/ml RNase A in Tris-EDTA [TE] buffer [10 mM Tris and 1 mM EDTA]) and treated at 37°C for 40 min. DNA cleanup was performed using SPRI beads (Ampure XP, Beckman Coulter) to enrich small DNA fragments from the Mnase DNA cuts and remove large DNA fragments that might result from spontaneous DNA breaks. First, a reverse SPRI cleanup was performed by adding 0.8× (24 µL) SPRI beads, followed by 5 min incubation at RT. The supernatant was collected, and the remaining small DNA fragments were purified by adding additional 1× (30 µL) SPRI beads and 5.4× (162 µL) isopropanol and incubating 5 min at RT. The beads were washed twice with 85% EtOH, and finally, DNA was eluted in 30 µL of 0.1× TE buffer.

### ChEC-seq library preparation and sequencing

Library preparation was performed as described^77^, with modifications. Following RNase treatment and reverse SPRI cleanup, the DNA fragments served as an input to an end-repair and A-tailing (ERA) reaction. 5.4 µL ERA reaction was prepared (1× T4 DNA ligase buffer [NEB], 0.5 mM dNTPs, 0.25 mM ATP, 2.75% PEG 4000, 6U T4 PNK [NEB], 0.5U T4 DNA

Polymerase [Thermo Scientific], 0.5U Taq DNA polymerase [Bioline]) and added to 14.6 µL of each sample and incubated for 20 min at 12°C, 15 min at 37°C and 45 min at 58°C in a thermocycler. After the ERA reaction, reverse SPRI cleanup was performed by adding 0.5× (10 µL) SPRI beads (Ampure XP, Beckman Coulter). The supernatant was collected, and the remaining small DNA fragments were purified with additional 1.3× (26 µL) SPRI beads and 5.4× (108 µL) isopropanol. After washing with 85% EtOH, small fragments were eluted in 17 µL of 0.1× TE buffer; 16.4 µL elution was taken into 40 µL ligation reaction (1× Quick ligase buffer [NEB], 4000U Quick ligase [NEB], and 6.4 nM Y-shaped barcode adaptors with T-overhang and incubated for 15 min at 20°C in a thermocycler. After incubation, the ligation reaction was cleaned by performing a double SPRI cleanup: first, a regular 1.2× (48 µL) SPRI cleanup was performed and eluted in 30 µL 0.1× TE buffer. Then instead of separating the beads, an additional SPRI cleanup was performed by adding 1.3× (39 µL) HXN buffer (2.5 M NaCl, 20% PEG 8000) and final elution in 24 µL 0.1× TE buffer; 23 µL elution were taken into 50 µL enrichment PCR reaction (1× Kappa HIFI [Roche], 0.32 µM barcoded Fwd primer and 0.32 µM barcoded Rev primer) and incubated for 45 s in 98°C, 16 cycles of 15s in 98°C and 15s in 60°C, and a final elongation step of 1 min at 72°C in a thermocycler.

The final libraries were cleaned using 1.1× (55 µL) SPRI and eluted in 15 µL 0.1× TE buffer. Library concentration and size distribution were quantified by Qbit (Thermo Scientific) and TapeStation (Agilent), respectively. For multiplexed next-generation sequencing (NGS), all barcoded libraries were pooled in equal amounts, and the final pool was diluted to 2 nM and sequenced on NovaSeq 6000/ NovaSeq X (Illumina). Sequence parameters were Read1: 61 nucleotides (nt), Index1: 8 nt, Index2: 8 nt, Read2: 61 nt.

### Co-Immunoprecipitation

NMuMG cells were plated at 65% confluency under standard conditions in 10cm dishes. Following 24 hours of culture, cells were added with the indicated ligands and incubated at 37°C for 45 min. Cells were washed twice with PBS, and lysed in Cellytic M lysis reagent (Sigma-Aldrich) added with 1:10 Phosphatase inhibitor (PhosSTOP, Sigma-Aldrich) and 1:400 Protease inhibitor Cocktail III (Sigma-Aldrich). Next, cells were incubated 15 minutes on ice, followed by the addition of Benzonase nuclease (1:1000) and incubation for 10 minutes at 30°C. Lysates were cleared by centrifugation (16,000g, 10 min, 4°C) to remove cell debris. Protein concentration was assessed using a BC Assay Protein Quantitation Kit (Interchim). 500ug of cleared lysates were then incubated on ice with antibodies: rabbit anti-SMAD2/3 (CST, #8685) or rabbit IgG (Abcam, #ab172730) which served as an isotype control. Following 1 hour of incubation, Protein A/G MagBeads (GenScript) was added and the mixture was incubated overnight at 4°C with head-to-tail rotation, followed by three 1 ml washes with a chilled TBS buffer. The target conjugates were then eluted from beads by mixing 4X LDS Sample Buffer (GenScript) with reducing agents (30mg/ml DTT and denaturant 4M Urea) and heating for 2 min at 95 °C followed by denaturing western blot.

### Western blot

Protein concentration was assessed using a BC Assay Protein Quantitation Kit (Interchim). 10 μg of total protein served as input and 20% of the immuno-precipitated proteins served as the pull-down fraction, were separated by SDS–PAGE on a 4–20% gradient SurePAGE™, Bis-Tris gel (GenScript). Gels were transferred onto nitrocellulose membranes using an iBlot3 Gel Transfer Device (Thermo Fisher Scientific). The membranes were blocked in 5% milk prepared in TBS–0.1% Tween and incubated in primary antibodies overnight at 4 °C, followed by washing and incubation with secondary antibody. Blots were developed using SuperSignal™

West Pico PLUS Chemiluminescent Substrate (Thermo Fisher Scientific) and the ChemiDoc XRS^+^ Imaging System (Bio-Rad). Band intensities were quantified with ImageJ analyzer software. Primary antibodies used: anti-SMAD1 (CST, #6944), anti-SMAD2/3 (CST, #8685), anti-SMAD4 (CST, #46535), and anti-Vinculin (Abcam, #ab129002) (diluted in 3% BSA in 0.1% Tween in TBS). Secondary antibodies used: Clean-Blot IP detection reagent HRP (Thermo Fisher Scientific) or Gout anti Rabbit HRP (Jackson labs). All primary antibodies were used in 1:1000 ratio, while secondary antibodies were used in 1:5000 ratio.

### Data Analysis

#### Flow cytometry

Initial analysis and gating of data was done using EasyFlow code for MATLAB. Further analysis was done using custom code in Python.

#### Batch correction

For every experiment, data sets from each repeat underwent batch correction in order to account for variations. We developed an orthogonal distance regression algorithm, which finds a straight line in the shape of

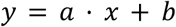

and minimizes its orthogonal distance (residues) to the two data sets, [x_i_] and [y_i_] with respect to the line parameters a (slope) and b (intercept) :

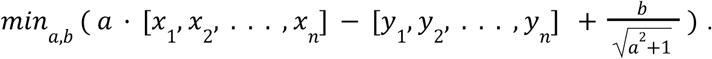

The resulting parameters describe the relative normalization (a) and offset (b) between the two data sets. For each experiment, we rescaled all data sets to one reference set via their specific fit parameters, thus correcting for the batch variations between them:

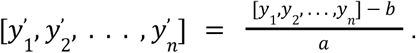

#### Hill curve fit

We created single ligand dose-response curves in order to extract key biochemical parameters specific to each ligand. The responses (R_i_) in dependence of the ligand concentrations (c_i_) were fitted to a Hill curve of the following shape:

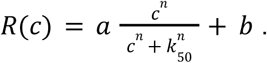

Its parameters denote the maximum response (a), the concentration resulting in half of that maximum response or EC_50_ (k_50_), and the offset (b).

### ChEC-Seq

#### ChEC-seq NGS data processing

Raw reads from ChEC-seq libraries were demultiplexed using bcl2fastq (Illumina), and adaptor dimers and short reads were filtered out using cutadapt^78^ with parameters: “—O 10 –pair-filter = any –max-n 0.8 –action = mask”. Filtered reads were subsequently aligned to the mm10 genome using Bowtie2^79^ with the options “--end-to-end --trim-to 40 --very-sensitive”.

#### Peak Calling and Filtering

For ChEC-seq experiments, peak calling was performed using MACS3^81^ with parameters --format BAMPE -g mm for mm10, along with options --nonmodel –no-lambda --keep-dup all. The called peaks were subsequently filtered using a custom Python script to remove peaks overlapping blacklisted regions. The peaks were then combined between the replicates using the IDR procedure with ChIP-R^82^. To generate the consensus set of peaks for further analysis, the two (mix and BMP4 addition) peak sets were merged and resized to a 600 bp window centered on the average midpoint or the initial summit of the overlapping and non-overlapping peaks, respectively, using the GenomicRanges library in R.

#### De novo motif analysis

De novo motif analysis was performed using HOMER (findMotifsGenome.pl) using the default parameters.

#### BigWig Normalization and Averaging

BigWig coverage files were generated from BAM alignments using deepTools^83^ bamCoverage, normalizing by Counts Per Million (CPM). The signal between repeats was then averaged using bigwigAverage.

#### Differential peak analysis and de novo motif finding

The sum of reads per peak in the unified set of peaks was calculated (see Peak Calling and Filtering) for both conditions using the respective BigWig files. And the enhanced-binding peaks for each condition were defined as those with log_2_(FC) > 1 between the calculated sums.

#### Differential k-mer enrichment analysis

To identify short sequences associated with condition-specific SMAD1 binding, we performed differential 7-mer enrichment analysis using the monaLisa Bioconductor package. Sequences from BMP-enhanced and mix-enhanced SMAD1 binding sites were extracted from the mm10 genome assembly. K-mer enrichment was computed using calcBinnedKmerEnr with k = 7, collapsing reverse complements, and using a genome-matched background nucleotide model. For each 7-mer, a log₂ fold-change was calculated as the difference in enrichment between the two condition-bins, and statistical significance was taken as the maximum −log_10_(adjusted p-value) across the two bins. Significantly enriched 7-mers were defined as those exceeding a −log_10_ (adjusted p-value) threshold of 4.

#### Average signal at motif sites

To examine SMAD1 binding patterns at predicted motifs, we computed average ChEC-seq signal profiles centered on motif occurrences within differential binding sites. Motif positions were identified by scanning the combined set of differential SMAD1 peaks against the indicated motifs (Fig. 5F) using matchMotifs (motifmatchr) with a genome-wide nucleotide background model. For each motif, the average signal from BMP, mix, and Control (no ligand addition) conditions’ average BigWig files was extracted over a 400-bp window centered on the motif midpoint, with minus-strand instances reverse-complemented to maintain consistent orientation. The mean signal across all sites was computed at each position, resulting in average binding profiles for each motif.

### Processing of RNA-sequencing data

#### MARS-seq analysis pipeline

RNA-seq data generated using the MARS-seq protocol were processed and analyzed via the UTAP pipeline. Raw FASTQ files were first quality-checked and trimmed using Cutadapt. Reads were aligned to the mouse genome (mm10) using STAR. Gene expression quantification was performed by counting uniquely mapped reads per gene using HTSeq-count, guided by the Gencode M25 annotation. For unique molecular identifier (UMI)-based deduplication, the MARS-seq analysis incorporated UMI collapsing at the gene level to ensure accurate molecule counts. Normalization and differential expression analysis were carried out using DESeq2. Genes with adjusted p-values (Benjamini-Hochberg FDR) < 0.05 were considered significantly differentially expressed.

#### Modeling of transcriptional crosstalk using a factorial design

To rigorously define the nature of transcriptional crosstalk between the TGFβ and BMP signaling pathways, differential expression was modeled using a 2×2 factorial design within the DESeq2 framework (implemented via PyDESeq2 in Python). We used generalized linear model (GLM) to fit the data to a model with an interaction term between TGFβ and BMP.

#### Computation of directional shift metric

We consider genes that respond to a primary ligand only, and want to quantify the synergistic effect of a secondary ligand. Using the GLM we can extract the fold-change associated with each ligand and define:

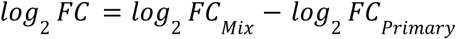

which captures the combined effect of the secondary ligand and the interaction term. Importantly, since these genes respond only to the primary ligand, the contribution of the secondary can be ignored.

Next, we wanted to capture whether this effect enhances or inhibits the original effect of the primary ligand. Specifically, if the primary ligand increases gene expression, then a positive *log*_2_ *FC* reflects an enhancement of the effect. However, if the primary ligand reduces gene expression, a positive *log*_2_ *FC* reflects an inhibition of the effect. To account for this we define:

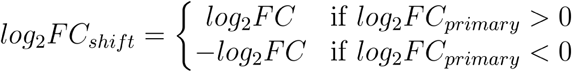

#### Statistical evaluation of transcriptional crosstalk

Statistical significance for these specific synergistic enhancements was determined natively within the DESeq2 model, testing against the null hypothesis that the combined condition is indistinguishable from the single-ligand condition. Raw p-values were adjusted using the Benjamini-Hochberg False Discovery Rate (FDR) procedure within the defined subsets of ligand-responsive genes.

#### Statistical evaluation of global enhancement via Monte-Carlo simulation

To assess whether the change in the overall distribution of log_2_FC_shift_ is statistically significant, we generated a null distribution using 10,000 Monte-Carlo simulations, by randomly sampling genes from the entire dataset while matching the sample size of the experimental gene set. The observed median shift was then compared to this distribution to compute empirical p-values.

#### EMT-score calculation

A set of 92 EMT- and MET-related genes (Table S2) was selected based on previously published EMT signatures. PCA was performed using the scikit-learn Python package (PCA module) on the gene expression matrix of the selected genes. The first principal component (PC1), which explained the greatest variance across conditions, was used as a composite EMT-score. Classical mesenchymal markers (e.g., *Snai1*, *Vim*, *Zeb1*) loaded positively on PC1, while epithelial markers (e.g., *Cdh1*, *Klf5*) loaded negatively.

## Acknowledgment

This work was done with critical advice from Dr. Hadas Keren-Shaul from the Genomics Sandbox unit at the Life Science Core Facility of Weizmann Institute of Science.

Y.E.A is supported by the Israel Science Foundation (grant 1105/20) and by a research grant from the Sygnet Fund. This research was supported by the Ilse Katz Institute for Materials Sciences and Magnetic Resonance Research.

## Declaration of interests

Y.E.A. is a scientific advisory board member and consultant of TeraCyte.

## Data availability

Bulk RNA sequencing data have been deposited at GEO: GSE254418 and are publicly available as of the date of publication.

## Notes

https://www.ncbi.nlm.nih.gov/geo/query/acc.cgi?acc=GSE254418

