## Supplemental Data 1 for "Asymmetric crosstalk between the BMP and TGFβ pathways resolves signaling ambiguity"

### Supplemental Information for: **Asymmetric crosstalk between the BMP and TGFβ pathways resolves signaling ambiguity**

#### Supplemental figures and tables

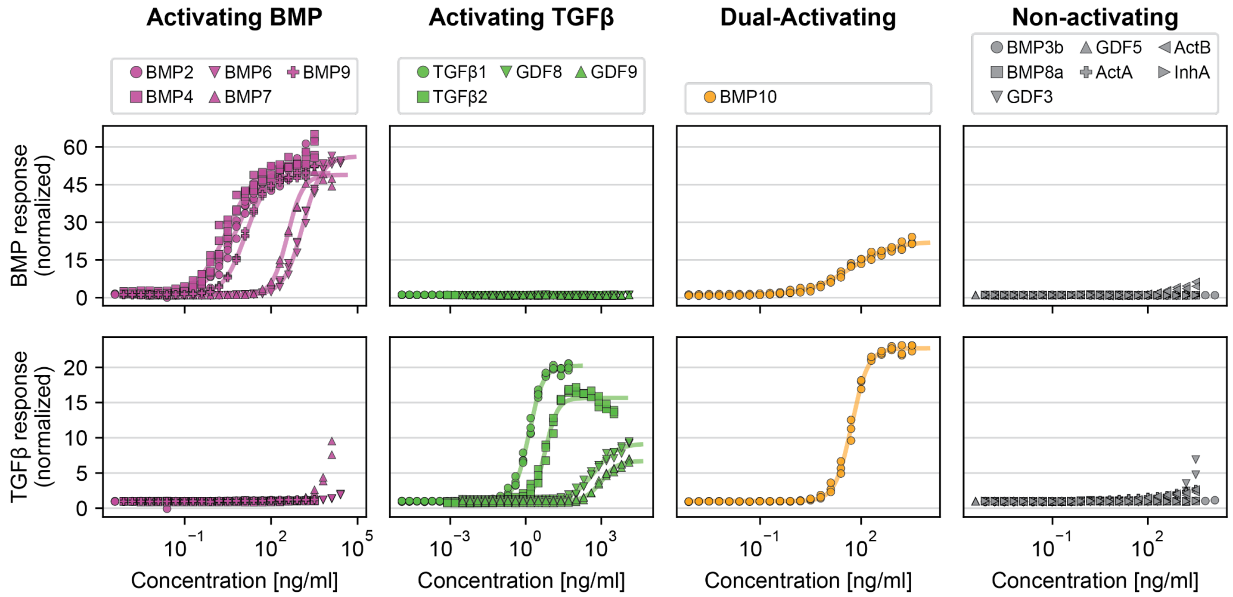


**Figure S1. Single ligand dose-response curves, related to Figure 1.** NMuMG cells with dual fluorescent reporters were incubated for 24h with a large set of BMP or TGFβ ligands. Each ligand was supplied at multiple concentrations across its entire dynamic range with 2-fold dilutions. Dose responses are plotted for ligands activating only a BMP response (purple), only a TGFβ response (green), dual response (orange), and no response (gray). The data shows two to four experimental repeats for each ligand. Responses were linearly corrected, fitted to a Hill function, and normalized to the baseline. Concentrations used and extracted Hill parameters are specified in Table S1.

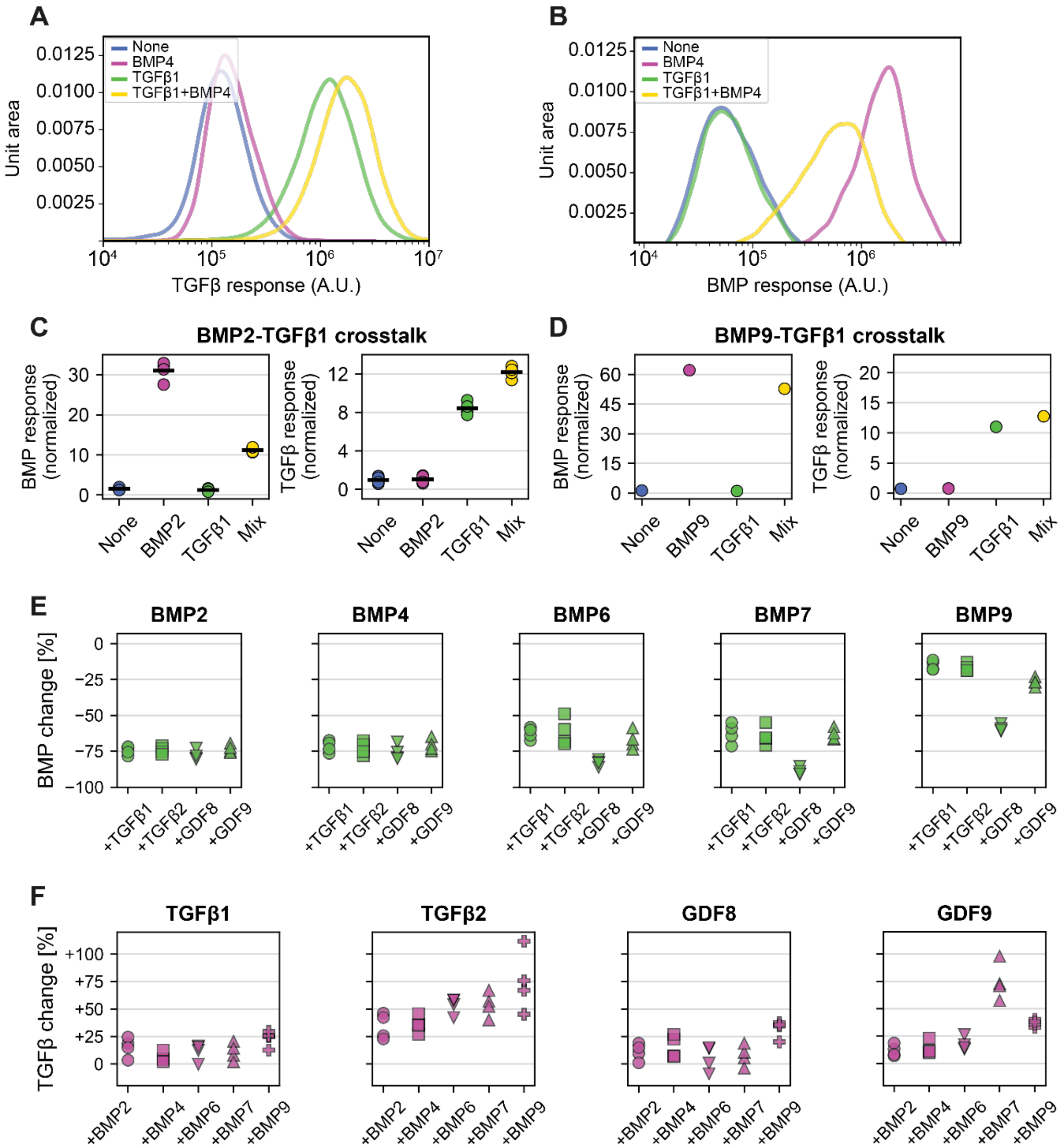


**Figure S2. Asymmetric crosstalk between BMP and TGFβ is observed across ligands and cell types, related to Figure 2.** (A-B) NMuMG dual reporter cells were cultured for 24h under four conditions: no ligands added (None), BMP4 at 31.25 ng/ml, TGFβ1 at 1.92 ng/ml, or a mix of both. (A) Histograms showing population-level mCitrine fluorescence, indicating TGFβ pathway activation. (B) Histograms showing population-level mCherry fluorescence, indicating BMP pathway activation. (C-D) NMuMG dual reporter cells were cultured for 24h under four conditions: no ligands added (None), a BMP ligand only, a TGFβ ligand only, or a mix of BMP and TGFβ ligands (Mix) (cf. Figure 2A). (C) 31.25 ng/ml of BMP2 and 1.92 ng/ml of TGFβ1 were used. The data shows n=3 repeats. (D) 33.33 ng/ml of BMP9 and 1.92 ng/ml of TGFβ1 were used. The data shows a single repeat. (E) Percent response change for BMP was calculated for BMP2, BMP4, BMP6, BMP7, and BMP9, when added with each of the TGFβ activating ligands. (F) Percent response change for TGFβ was calculated for TGFβ1, TGFβ2, GDF8, and GDF9, when added with each of the BMP activating ligands.

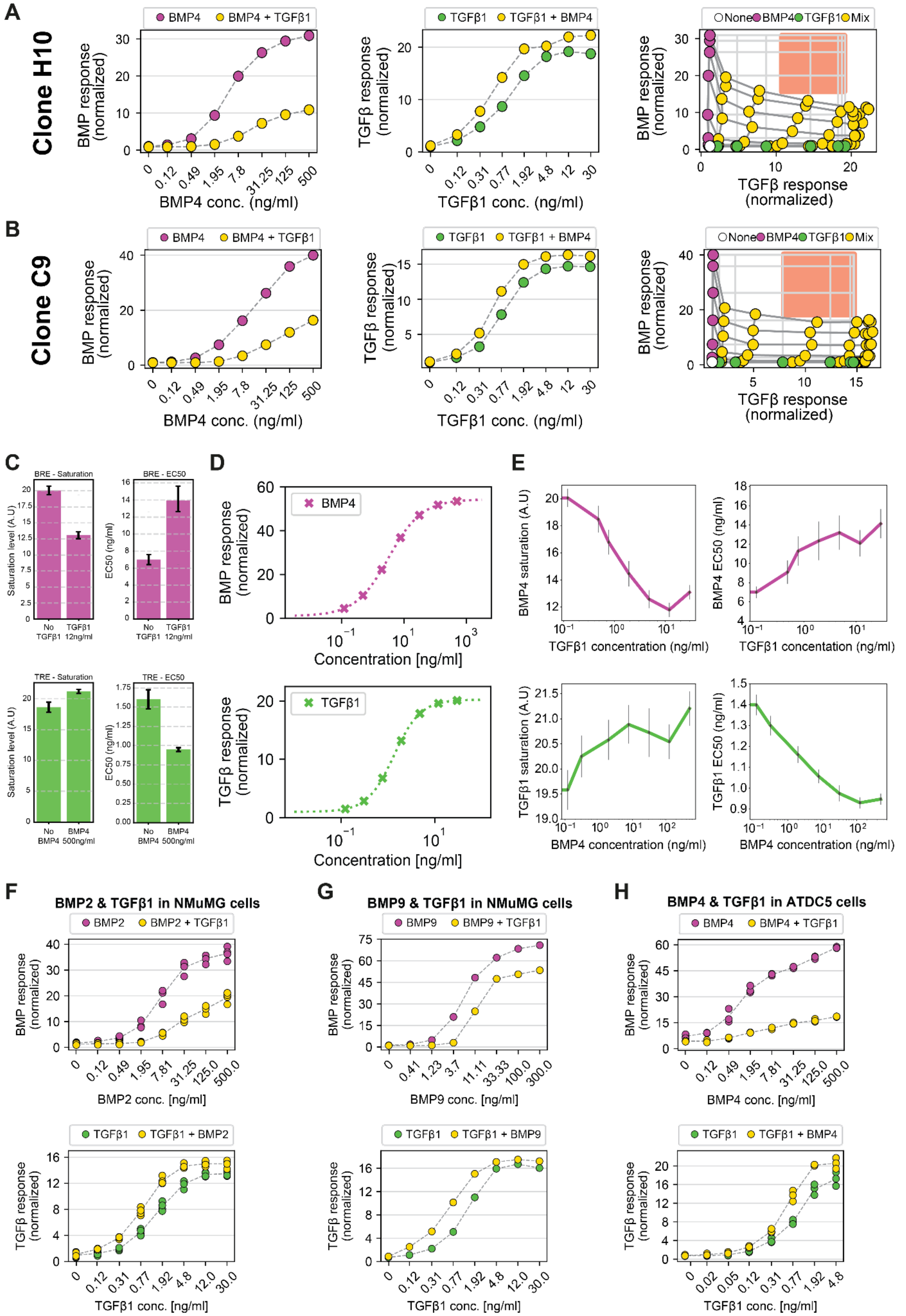
A
Clone H10
BMP4
BMP4 + TGFβ1
BMP response (normalized)
BMP4 conc. (ng/ml)
TGFβ1
TGFβ1 + BMP4
TGFβ response (normalized)
TGFβ1 conc. (ng/ml)
None BMP4 TGFβ1 Mix
TGFβ response (normalized)
B
Clone C9
C
BRE - Saturation
BRE - EC50
Saturation level (A.U)
EC50 (ng/ml)
No TGFβ1
TGFβ1 12ng/ml
TRE - Saturation
TRE - EC50
No BMP4
BMP4 500ng/ml
D
Concentration [ng/ml]
E
BMP4 saturation (A.U)
BMP4 EC50 (ng/ml)
TGFβ1 concentration (ng/ml)
TGFβ1 saturation (A.U)
TGFβ1 EC50 (ng/ml)
BMP4 concentration (ng/ml)
F
BMP2 & TGFβ1 in NMuMG cells
BMP2
BMP2 + TGFβ1
BMP2 conc. [ng/ml]
TGFβ1 + BMP2
TGFβ1 conc. [ng/ml]
G
BMP9 & TGFβ1 in NMuMG cells
BMP9
BMP9 + TGFβ1
BMP9 conc. [ng/ml]
TGFβ1 + BMP9
H
BMP4 & TGFβ1 in ATDC5 cells
BMP4 conc. [ng/ml]

**Figure S3. Asymmetric crosstalk reduces ambiguity across multiple ligand pairs and cell lines, related to Figure 2.** (A-B) Dose-response curves and two-dimensional concentration gradient assays of two additional clones of NMuMG dual reporter cells. Cf. Figures 2D-F. (C) Saturation and $EC_{50}$ levels extracted from the dose response of BMP (top) or TGFβ (bottom) in the presence or absence of the other ligand by fitting with a Hill function. (D) Detailed depiction of the concentration sampling for the 8x8 2D-gradients. Based on the Hill function generated by the calibration curves for BMP4 and TGFβ1, we chose near-saturating concentrations and six lower concentrations at log-equidistant intervals, as well as an eighth condition with no added ligands. (E) Change in the saturation level and $EC_{50}$ concentration of BMP4 (top) and TGFβ1 (bottom) over concentrations of the other ligand. (F-H) Dose-response curves of multiple ligand pairs and different cell types. Cf. Figures 2D,E (F) BMP2 and TGFβ1 in NMuMG dual reporter cells. TGFβ1 concentration in the BMP2 mixed dose-response curve is 30 ng/ml. BMP2 concentration in the TGFβ1 mixed dose-response curve is 500 ng/ml. Data shown contains n=4 biological repeats. (G) BMP9 and TGFβ1 in NMuMG dual reporter cells. TGFβ1 concentration in the BMP9 mixed dose-response curve is 30 ng/ml. BMP9 concentration in the TGFβ1 mixed dose-response curve is 300 ng/ml. Data shown contains n=1 biological repeats. (H) BMP4 and TGFβ1 in ATDC5 dual reporter cells. TGFβ1 concentration in the BMP4 mixed dose-response curve is 30 ng/ml. BMP4 concentration in the TGFβ1 mixed dose-response curve is 500 ng/ml. Data shown contains n=3 biological repeats.

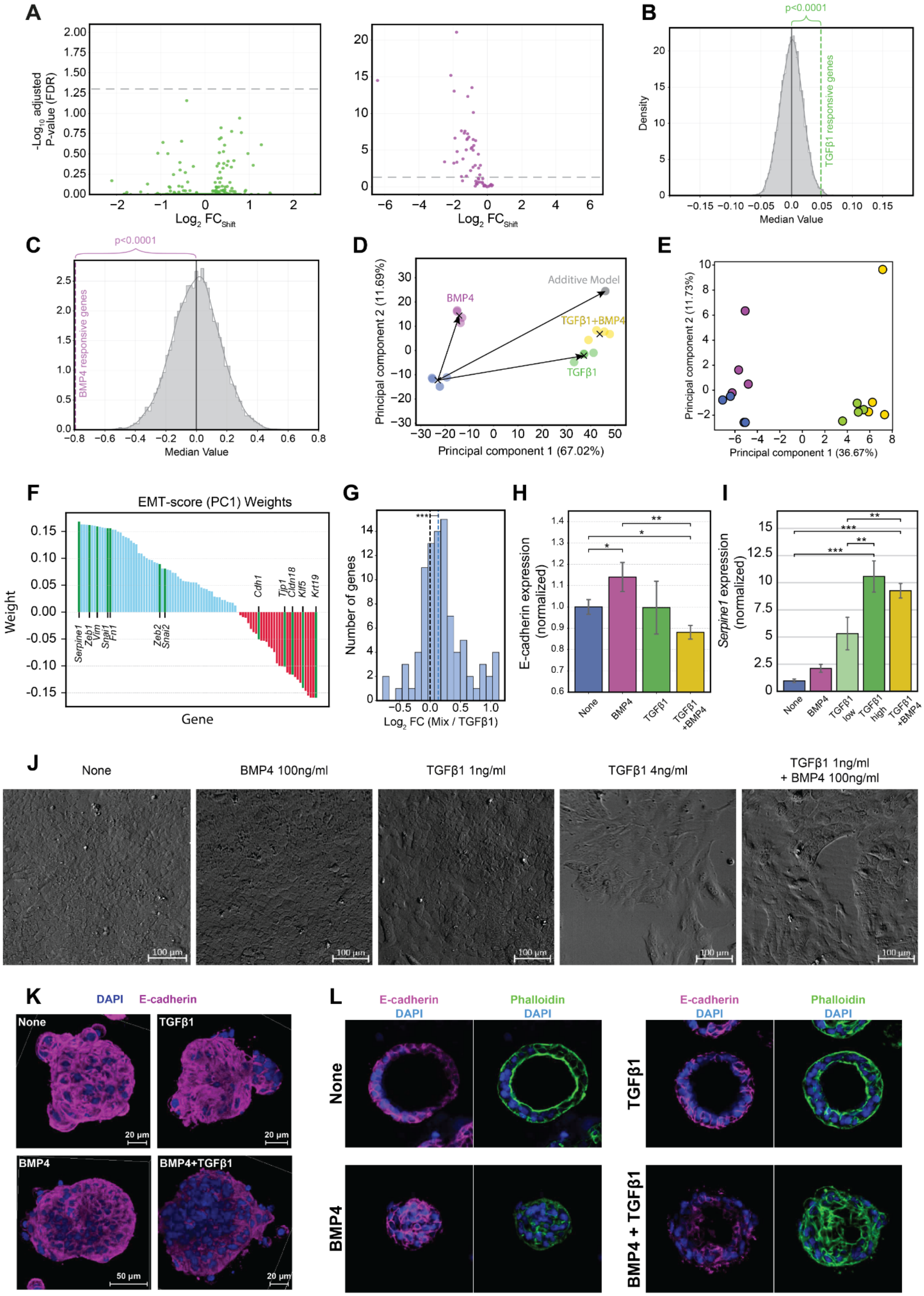

A
-Log10 adjusted P-value (FDR)
Log2 FCShift
B
p<0.0001
TGFβ1 responsive genes
Density
Median Value
C
p<0.0001
BMP4 responsive genes
Median Value
D
Additive Model
BMP4
TGFβ1+BMP4
TGFβ1
Principal component 2 (11.69%)
Principal component 1 (67.02%)
E
Principal component 2 (11.73%)
Principal component 1 (36.67%)
F
EMT-score (PC1) Weights
Weight
Gene
Serpine1
Zeb1
Vim
Snai1
Fn1
Zeb2
Snai2
Cdh1
Tjp1
Cldn18
Klf5
Krt19
G
Number of genes
Log2 FC (Mix / TGFβ1)
H
E-cadherin expression (normalized)
None
BMP4
TGFβ1
TGFβ1 +BMP4
I
Serpine1 expression (normalized)
None
BMP4
TGFβ1 low
TGFβ1 high
TGFβ1 +BMP4
J
None
BMP4 100ng/ml
TGFβ1 1ng/ml
TGFβ1 4ng/ml
TGFβ1 1ng/ml + BMP4 100ng/ml
100 μm
K
DAPI
E-cadherin
None
TGFβ1
BMP4
BMP4+TGFβ1
20 μm
50 μm
L
E-cadherin
DAPI
Phalloidin
DAPI
None
BMP4
TGFβ1
BMP4 + TGFβ1

**Figure S4. Asymmetric BMP-TGFβ crosstalk reshapes transcriptional programs and EMT signatures, related to Figure 3.** (A) Volcano plots showing deviation from an additive expectation under mixed stimulation for TGFβ1-responsive genes (left, green) and BMP4-responsive genes (right, purple). For each gene, the x-axis shows the directional shift relative to an additive model ($Log_2\,FC_{Shift}$), and the y-axis shows -$Log_{10}$ (adjusted p-value). Dashed gray line marks adjusted p-value (FDR) = 0.05. (B-C) Monte-Carlo significance test for global shifts in gene expression under BMP4 + TGFβ1 co-stimulation. Gray histograms show the null distribution of median $Log_2\,FC_{Shift}$ values obtained from randomly sampled gene sets; dashed line indicates the observed median for TGFβ1-responsive genes (B) or BMP4-responsive genes (C), with the corresponding p-value indicated. (D) PCA of all responsive genes across four conditions: no ligand added (blue), TGFβ1 (green), BMP4 (purple), and a combination of both (yellow). Crosses (x) denote group means. The additive model prediction (gray circle) is shown for comparison with the measured mixed condition. (E) PCA of curated EMT- and MET-marker genes across conditions. Individual biological replicates are shown for Control (blue), BMP4 (purple), TGFβ1 (green), and Mix (yellow) conditions. (F) EMT score was defined by the weights of the first principal component (PC1) from the PCA performed on the EMT- and MET-marker genes. Genes with positive contributions to the EMT score are shown in blue, while negative contributors are shown in red. Representative EMT and MET hallmark genes are labeled. (G) Distribution of EMT/MET gene expression changes in the mixed condition relative to TGFβ1 alone ($Log_2\,FC$ Mix/TGFβ1). The black dashed vertical line represents a $Log_2\,FC$ of 0, while the blue dashed vertical line marks the median of the observed distribution. (H) Flow cytometry analysis of E-cadherin expression levels in NMuMG cells following 36h stimulation under treatment with no ligand (blue), 100 ng/ml of BMP4 (purple), 1 ng/ml of TGFβ1, or a combination of both ligands. Data represents the normalized median fluorescence intensity, calculated relative to the mean of the no-ligand control group. Bar graphs show the mean ± s.d. of n=3 replicates. (I) qPCR of *Serpine1* mRNA following treatment with TGFβ1 (low: 1 ng/ml; high: 4 ng/ml) with or without BMP4 (100 ng/ml). Bar graphs show mean ± s.d. of n=2 biological replicates. (J) Microscopy brightfield images showing the morphological transition of NMuMG cells after 36h of treatment with the indicated concentrations of BMP4, TGFβ1, or the combination of both. Scale bars = 100 μm. (K) 3D confocal images of breast organoids immunostained for E-cadherin (magenta) and DAPI (blue), showing responses to four treatment conditions: no ligand added (None), TGFβ1 at 2 ng/ml, BMP4 at 100 ng/ml, and a combination of both (TGFβ1+BMP4). (L) Confocal slices of organoid cross-sections stained for E-cadherin (magenta), phalloidin (green), and DAPI (blue). In all figures, *P ≤ 0.05, **P ≤ 0.01, ***P ≤ 0.001, ****P ≤ 0.0001 using two-sided t-test.

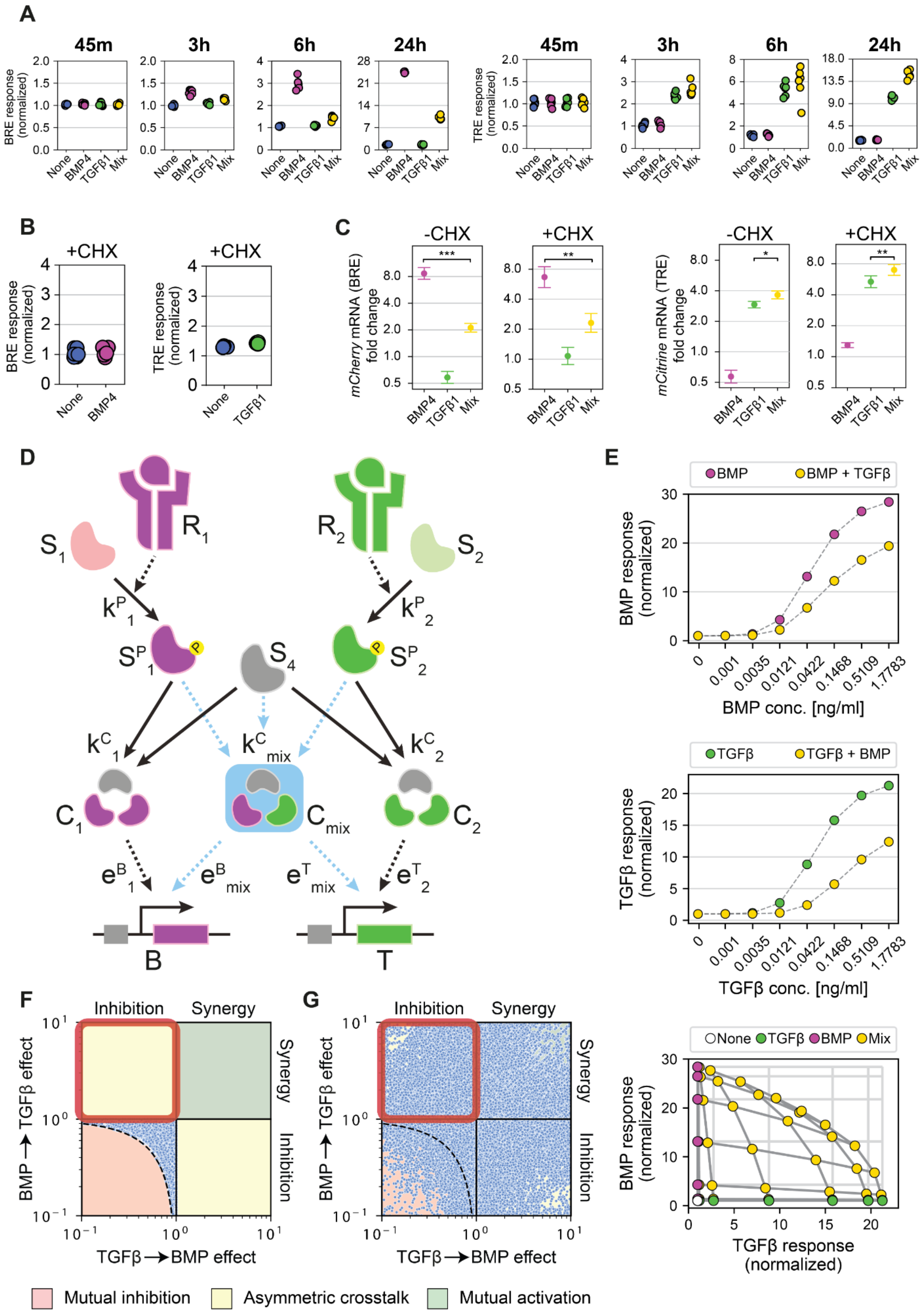

A
45m
3h
6h
24h
BRE response (normalized)
TRE response (normalized)
None
BMP4
TGFβ1
Mix
B
+CHX
C
-CHX
+CHX
mCherry mRNA (BRE) fold change
mCitrine mRNA (TRE) fold change
D
S1
R1
R2
S2
S4
C1
Cmix
C2
B
T
E
BMP
BMP + TGFβ
BMP response (normalized)
BMP conc. [ng/ml]
TGFβ
TGFβ + BMP
TGFβ response (normalized)
TGFβ conc. [ng/ml]
None
TGFβ
BMP
Mix
F
G
Inhibition
Synergy
BMP ⟶ TGFβ effect
TGFβ ⟶ BMP effect
Mutual inhibition
Asymmetric crosstalk
Mutual activation

**Figure S5. Mixed SMAD complexes expand the computational capacities of cellular signaling beyond canonical mutual antagonism, related to Figure 4.** (A) Dual-reporter NMuMG cells were treated with either no ligand (None, blue), BMP4 at 31.25 ng/ml (purple), TGFβ1 at 1.92 ng/ml (green), or both ligands combined (Mix, yellow) for 45 minutes, 3, 6, or 24 hours. mCherry (BRE, left) and mCitrine (TRE, right) fluorescence levels were quantified by flow cytometry to assess temporal signaling dynamics. (B, C) To test the role of feedback regulation, dual-reporter NMuMG cells were treated with or without cycloheximide (CHX) during a 6-hour exposure to the same four ligand conditions described above. (B) Flow cytometry was used to measure mCherry (BRE, left) and mCitrine (TRE, right) accumulation. (C) qPCR analysis was performed to quantify mCherry (BRE, left) and mCitrine (TRE, right) transcript levels, relative to the no-ligand control, in the presence (+CHX, right) or absence (-CHX, left) of cycloheximide. (D) Schematic depiction of the intracellular model. Ligand receptor complexes ($R_i$) control the phosphorylation rate ($k^P{}_i$) of SMAD proteins ($S_i$). The phosphorylated SMADs ($S^P{}_i$) combine with co-SMAD4 ($S_4$) to form homogeneous complexes ($C_i$) or mixed complexes ($C_{mix}$). The complexes induce a transcriptional response for a BMP target gene (B) or a TGFβ target gene (T) according to their efficacies ($e^B{}_i$, $e^T{}_i$, $e^B{}_{mix}$, $e^T{}_{mix}$). (E) Modeling results of simulating a two-dimensional concentration gradient assay without mixed complexes ($k^C{}_{mix} = 0$). Cf. Figures 2D,E. (F-G) Parameter scan of the intracellular model without mixed complexes ($k^C{}_{mix} = 0$) (F) or with mixed complexes ($k^C{}_{mix} > 0$) (G). For each data point (blue), four conditions (no ligands added, BMP ligand only, TGFβ ligand only, and a mix of both ligands) were simulated. We then calculated, for both pathways, the percent response change between the mixed condition and the respective single ligand condition. The quadrant colors indicate the type of crosstalk: mutual inhibition (red), asymmetric (yellow), and mutual activation (green). The red square marks the asymmetric crosstalk regime that corresponds to our experimental observations, where the addition of TGFβ decreases the BMP response, while BMP addition enhances the TGFβ response. For detailed parameter scan range and sampling steps, see Methods section. In all figures, $*P \leq 0.05$, $**P \leq 0.01$, $***P \leq 0.001$, $****P \leq 0.0001$, determined using a paired t-test.

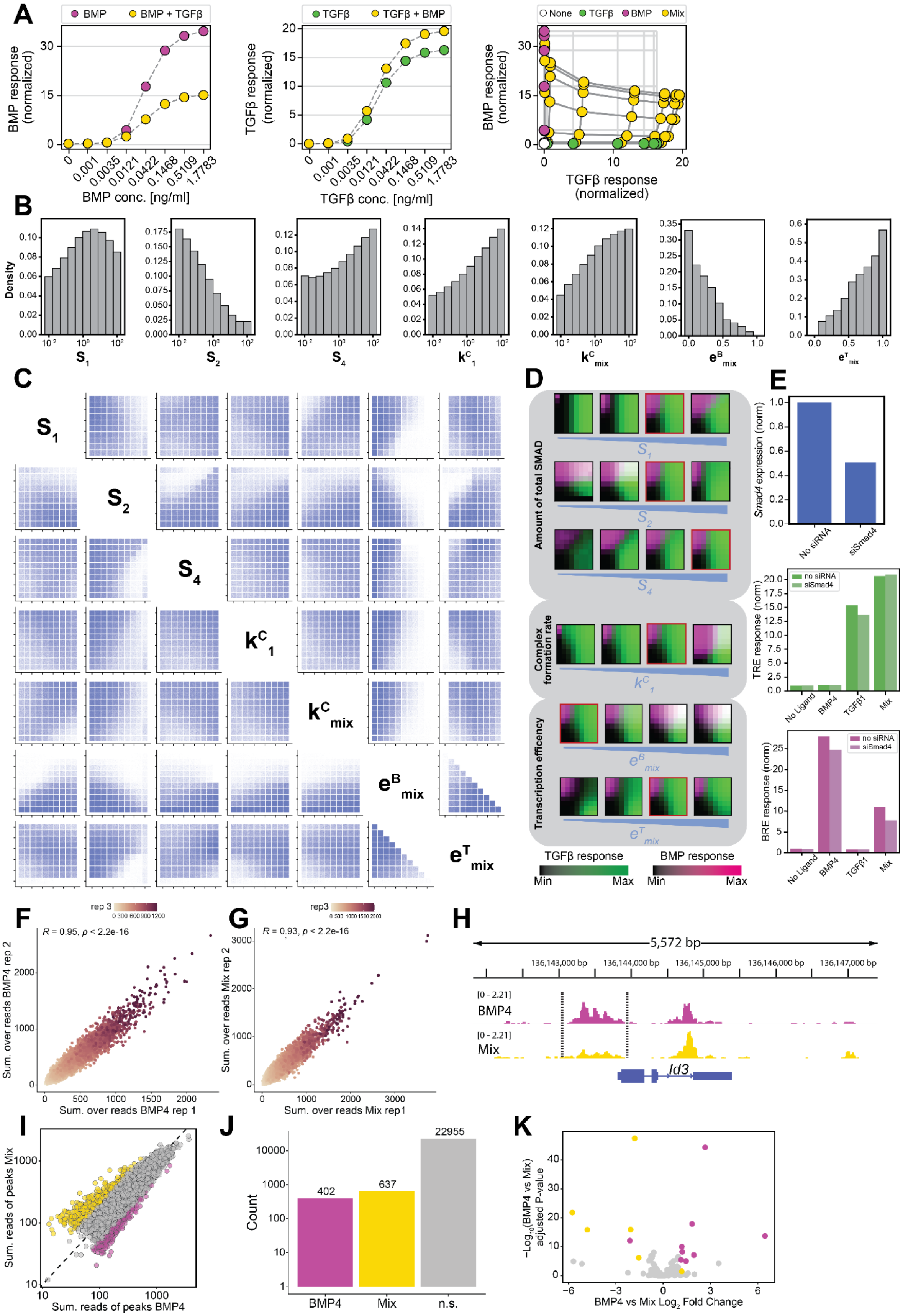

A
BMP
BMP + TGFβ
BMP response (normalized)
BMP conc. [ng/ml]
TGFβ
TGFβ + BMP
TGFβ response (normalized)
TGFβ conc. [ng/ml]
None
Mix
TGFβ response (normalized)
B
Density
C
D
Amount of total SMAD
Complex formation rate
Transcription efficency
TGFβ response
BMP response
Min
Max
E
Smad4 expression (norm)
No siRNA
siSmad4
TRE response (norm)
no siRNA
No Ligand
BMP4
TGFβ1
BRE response (norm)
F
R = 0.95, p < 2.2e-16
Sum. over reads BMP4 rep 2
Sum. over reads BMP4 rep 1
G
R = 0.93, p < 2.2e-16
Sum. over reads Mix rep 2
Sum. over reads Mix rep1
H
5,572 bp
136,143,000 bp
136,144,000 bp
136,145,000 bp
136,146,000 bp
136,147,000 bp
[0 - 2.21]
Id3
I
Sum. reads of peaks Mix
Sum. reads of peaks BMP4
J
Count
402
637
22955
n.s.
K
BMP4 vs Mix Log2 Fold Change

**Figure S6. Asymmetric BMP–TGFβ crosstalk is robust and emerges across diverse parameter spaces, related to Figure 5.** (A) Simulations of the model incorporating nonlinear transcriptional response across concentration gradients of BMP (left) or TGFβ (middle) ligands (purple or green, respectively), compared to the same gradients together with a constant ligand from the opposing pathway (yellow), as well as a full two-dimensional ligand gradient with BMP and TGFβ ligands. The BMP response and TGFβ response are plotted for all combinatorial conditions, including no stimulation (white), BMP ligand only (purple), TGFβ only (green), and mixed conditions (yellow). (B,C) A comprehensive parameter scan of our mathematical model was conducted by sampling 8 values for each parameter. Simulations were filtered to retain only those that reproduced the experimentally observed asymmetric crosstalk, where TGFβ signaling dominates over BMP. (B) One-dimensional distributions show how individual parameter values are enriched in the subset of simulations that reproduce an asymmetric crosstalk. (C) Two-dimensional pairwise plots reveal relationships and dependencies between parameters within the selected subset. (D) To assess how individual parameters influence crosstalk behavior, we performed a series of simulations in which one parameter was varied while the others were held constant. Rows represent the effect of varying each parameter. Parameter sets that replicated the experimentally observed crosstalk (see Figure 4E–G) are outlined in red. (E) NMuMG dual reporter cells were transfected with siRNA against SMAD4 for 24h (top, blue) and further cultured for an additional 24h under four conditions: no ligands added, BMP4 at 31.25 ng/ml, TGFβ1 at 1.92 ng/ml, or a mix of both. Pathway activation of TGFβ (TRE, green) and BMP (BRE, purple) in the presence or absence of siRNA-mediated knockdown of SMAD4 was measured using flow cytometry. ChEC-seq analysis of SMAD1 binding. (F-G) Scatter plots comparing the sum over reads of SMAD1 binding for n=3 replicates under stimulation by BMP4 only (48 ng/ml, F) and BMP4 + TGFβ1 (Mix; 1.92 ng/ml TGFβ1, G). Sum over reads are shown for repeats 1, 2, and 3 in the x-axis, y-axis, and color scale, respectively. Pearson correlation coefficients and P-values are provided for each comparison. (H) SMAD1-MNase occupancy at the *Id3* locus was measured following treatment with BMP4 only (purple) or with TGFβ1 (Mix, yellow). (I) Scatter plot comparing read density at genomic peaks between BMP4 only (48 ng/ml, purple) and BMP4 + TGFβ1 (Mix; 1.92 ng/ml TGFβ1, yellow) conditions. Peaks are classified based on differential occupancy: enhanced in BMP4 ($Log_2FC > 1$; purple), enhanced in Mix ($Log_2FC < -1$; yellow), or not significantly changed (gray). (J) Quantification of the peak categories identified in (I), showing the total count of BMP4-enhanced, Mix-enhanced, and non-significant (n.s.) peaks. (K) All genes associated with differential ChEC-seq peaks located near their promoter (-1000, +200 bp surrounding the TSS) were identified, and their differential mRNA expression between BMP4 and mixed (BMP4 + TGFβ1) conditions is shown as a volcano plot. Significantly differentially expressed genes are color-coded according to whether the nearby peak is stronger under BMP4 (purple) or mixed (yellow) stimulation, linking differential SMAD1 occupancy to transcriptional output.

| **Table S1. Single ligand calibration parameters.** | | | | | | | | | |
|---|---|---|---|---|---|---|---|---|---|
| **Ligand** | **Max conc. [ng/ml]** | **TRE Hill parameters** | | | | **BRE Hill parameters** | | | |
| | | **Max. *a* [a.u.]** | **Hill coef. *n*** | **$EC_{50}$ $k_{50}$ [ng/ml]** | **Offset *b* [a.u.]** | **Max *a* [a.u.]** | **Hill coef. *n*** | **$EC_{50}$ $k_{50}$ [ng/ml]** | **Offset *b* [a.u.]** |
| **TGFβ1** | 50 | 19.25 | 1.53 | 1.34 | 1 | - | - | - | - |
| **TGFβ2** | 3200 | 14.66 | 1.79 | 6.12 | 1 | - | - | - | - |
| **GDF8** | 12800 | 8.17 | 0.99 | 510 | 1 | - | - | - | - |
| **GDF9** | 12800 | 5.70 | 1.37 | 1003 | 1 | - | - | - | - |
| **BMP2** | 1200 | - | - | - | - | 52.17 | 0.86 | 6.32 | 1 |
| **BMP4** | 3200 | - | - | - | - | 53.38 | 0.81 | 3.25 | 1 |
| **BMP6** | 25600 | - | - | - | - | 55.31 | 1.30 | 1264 | 1 |
| **BMP7** | 12800 | - | - | - | - | 47.85 | 1.64 | 372.80 | 1 |
| **BMP9** | 3200 | - | - | - | - | 48.60 | 1.14 | 13.19 | 1 |
| **BMP10** | 3200 | 21.71 | 1.78 | 52.02 | 1 | 25.66 | 0.71 | 47.70 | 1 |
| **BMP3b** | 12800 | - | - | - | - | - | - | - | - |
| **BMP8a** | 3200 | - | - | - | - | - | - | - | - |
| **GDF3** | 3200 | - | - | - | - | - | - | - | - |
| **GDF5** | 1600 | - | - | - | - | - | - | - | - |
| **Activin-A** | 3200 | - | - | - | - | - | - | - | - |
| **Activin-B** | 3200 | - | - | - | - | - | - | - | - |
| **Inhibin-A** | 3200 | - | - | - | - | - | - | - | - |

**Table S2. EMT- and MET-related marker genes.**

| | | | |
|---|---|---|---|
| *Ablim1* | *Acta2* | *Acvrl1* | *Adam12* |
| *Anxa6* | *Atp10a* | *Bmi1* | *Cald1* |
| *Cdh1* | *Cdh2* | *Cldn18* | *Cldn4* |
| *Cldn9* | *Cmtm3* | *Col1a1* | *Col1a2* |
| *Col3a1* | *Col5a1* | *Col5a2* | *Col6a1* |
| *Col6a2* | *Col6a3* | *Dab2* | *Dsp* |
| *Emilin1* | *Epha1* | *Epn3* | *Erbb3* |
| *Esrp2* | *Evpl* | *Fam83b* | *Fn1* |
| *Fosl2* | *Foxa1* | *Foxc1* | *Foxc2* |
| *Foxf2* | *Fstl1* | *Fyn* | *Glt8d2* |
| *Gnai2* | *Grhl1* | *Grhl3* | *Hlx* |
| *Irf6* | *Jup* | *Klf5* | *Krt10* |
| *Krt14* | *Krt18* | *Krt19* | *Krt5* |
| *Krt6a* | *Krt7* | *Krt8* | *Lad1* |
| *Map7* | *Marveld3* | *Mmp2* | *Mmp9* |
| *Myc* | *Nid2* | *Olfml2b* | *Olfml3* |
| *Pcolce* | *Pdgfra* | *Pdgfrb* | *Perp* |
| *Pkp1* | *Pmp22* | *Postn* | *Ppl* |
| *Prrg4* | *Serpine1* | *Snai1* | *Snai2* |
| *Sox10* | *Sox4* | *Sparc* | *St3gal2* |
| *Syt11* | *Tcf4* | *Tead2* | *Timp2* |
| *Tjp1* | *Trim29* | *Twist1* | *Twist2* |
| *Vcan* | *Vim* | *Zeb1* | *Zeb2* |

| **Table S3. Simulation Parameters, related to Figure 4.** | | | |
|---|---|---|---|
| | **Fig. 4 (E-G)** | **Extended Data Fig. 5E** | **Extended Data Fig. 6A** |
| **Initial (free) amounts** | | | |
| **$S_1$** | 45 | 45 | 80 |
| **$S_2$** | 65 | 65 | 20 |
| **$S^P{}_1$** | 0 | 0 | 0 |
| **$S^P{}_2$** | 0 | 0 | 0 |
| **$S_4$** | 65 | 30 | 100 |
| **$C_1$** | 0 | 0 | 0 |
| **$C_2$** | 0 | 0 | 0 |
| **$C_{mix}$** | 0 | 0 | 0 |
| **Parameters** | | | |
| **$k^P{}_1$** | 1 | 1 | 1 |
| **$k^P{}_2$** | 1 | 1 | 1 |
| **$k^C{}_1$** | 0.25 | 0.25 | 1 |
| **$k^C{}_2$** | 0.25 | 0.25 | 1 |
| **$k^C{}_{mix}$** | 4.5 | 0 | 0.75 |
| **$e^B{}_1$** | 1 | 1 | 1 |
| **$e^B{}_2$** | 0 | 0 | 0 |
| **$e^B{}_{mix}$** | 0 | 0 | 0.125 |
| **$e^T{}_1$** | 0 | 0 | 0 |
| **$e^T{}_2$** | 1 | 1 | 1 |
| **$e^T{}_{mix}$** | 0.6 | 0 | 0.9 |

| **Table S4. Specification of ligands used, related to STAR Methods.** | | |
|---|---|---|
| **Ligand name** | **Vendor** | **cat#** |
| TGFβ1 | R&D Systems | 7666-MB-005 |
| TGFβ2 | R&D Systems | 7346-B2-005 |
| BMP2 | R&D Systems | 355-BM-010 |
| BMP3b | R&D Systems | 1543-BP-025 |
| BMP4 | R&D Systems | 5020-BP-010 |
| BMP6 | R&D Systems | 6325-BM-020 |
| BMP7 | R&D Systems | 5666-BP-010 |
| BMP8a | R&D Systems | 7540-BP-025 |
| BMP9 | R&D Systems | 5566-BP-010 |
| BMP10 | R&D Systems | 6038-BP-025 |
| GDF3 | R&D Systems | 9009-GD-010 |
| GDF5 | R&D Systems | 853-G5-050 |
| GDF8 | R&D Systems | 788-G8-010 |
| GDF9 | R&D Systems | 739-G9-010 |
| Activin-A | R&D Systems | 338-AC-010 |
| Activin-B | R&D Systems | 8260-AB-010 |
| Inhibin-A | R&D Systems | 8346-IN-010 |

| Table S5. Specification of primers used for quantitative PCR. | | |
|---|---|---|
| **Gene** | **Forward primer 5'→3'** | **Reverse primer 5'→3'** |
| *Sdha* | AGTGGGCTGTCTTCCTTAAC | GGATTGCTTCTGTTTGCTTGG |
| *mCitrine* | TTCAAGATCCGCCACAACAT | CTTCTCGTTGGGGTCTTTGC |
| *mCherry* | GAACGGCCACGAGTTCGAGA | CTTGGAGCCGTACATGAACTGAG |
| *Serpine1* | TCAATGACTGGGTGGAAAGGCA | AGGCGTGTCAGCTCGTCTAC |
| *Smad4* | CAAACCATCCAACACCCGCC | ATACTGGCCGGCTGACTTGT |

| Table S6. Model Variables. | |
|---|---|
| $S_1$, $S_2$, $S_4$ | SMAD proteins (free amounts): $S_1$ represents all SMAD proteins affiliated with the BMP pathway (SMAD1/5/8), $S_2$ represents all SMAD proteins affiliated with the TGFβ pathway (SMAD2/3), and $S_4$ represents SMAD4. |
| $S^P{}_1$, $S^P{}_2$ | phosphorylated SMAD proteins (pSMADs; free amounts):<br>Products of phosphorylation of $S_1$, $S_2$ at respective phosphorylation rates $k^P{}_{ij}$. $S_4$ is not phosphorylated. |
| $C_1$, $C_2$, $C_{mix}$ | trimeric SMAD complexes: $C_{ij}$ is composed of two phosphorylated SMADs ($S^P{}_i$, $S^P{}_j$) and one $S_4$. Complexes are formed at rates $k^C{}_1$. |
| B, T | transcriptional response: the trimeric SMAD complexes $C_{ij}$ induce target gene transcription response B, T at specific efficacies $e^B{}_{ij}$. |

| Table S7. Model Parameters. | |
|---|---|
| $R_1$, $R_2$ | amounts of ligand-receptor-complexes: each ligand binds to its specific pair of type I and type II receptor subunits. |
| $S^0{}_1$, $S^0{}_2$, $S^0{}_4$ | Initial amounts of SMAD proteins: $S_1$ represents all SMAD proteins affiliated with ligand 1 pathway (not distinguishable individually), $S_2$ respectively and $S_4$ is SMAD4. |
| $k^P{}_1$, $k^P{}_2$ | phosphorylation rates: $k^P{}_{ij}$ determines if and at what rate the ligand-receptor-complex $R_j$ phosphorylates SMAD protein $S_i$. |
| $k^C{}_1$, $k^C{}_2$, $k^C{}_{mix}$ | rates of SMAD-complex formation: $k^C{}_{ij}$ determines if and at what rate SMAD proteins $S_i$, $S_j$, and $S_4$ form a trimeric SMAD complex $C_{ij}$. |
| $e^B{}_1$, $e^B{}_{mix}$<br>$e^T{}_2$, $e^T{}_{mix}$ | efficacies of SMAD-complex induced transcription: $e^g{}_i$ determines if and at what rate SMAD complex $C_i$ binds to control the expression level of gene g. |

### Supplementary Information

#### Mathematical model

We analyze the intracellular layer of the BMP and TGFβ pathways using a mathematical model describing the biochemical binding-unbinding interactions (Extended Data Fig. 4a). The model describes the phosphorylation of SMADs, the formation of SMAD complexes and the induction of the transcriptional response. In particular, this model considers crosstalk between the pathways at these three levels.

#### SMAD phosphorylation

We start by considering the SMAD phosphorylation process. In this model, we use $S_1$ to represent all SMAD proteins affiliated with the BMP pathway (SMAD1/5/8) and $S_2$ to represent all SMAD proteins affiliated with the TGFβ pathway (SMAD2/3). The phosphorylation of $S_1$ and $S_2$ is controlled by the number of receptor complexes containing BMP ($R_1$) or TGFβ ($R_2$). We denote the phosphorylated proteins as $S^P{}_1$ and $S^P{}_2$. Our experimental results show no crosstalk at the level of SMAD phosphorylation (Fig. 3). Therefore, we consider each pathway to only phosphorylate its corresponding SMAD proteins. We denote the rates of $S_1$ and $S_2$ phosphorylation by $k^P{}_1$ and $k^P{}_2$, respectively. The steady-state equations for the phosphorylation process are given by (Eq. 1-2).

$$S^P{}_1 = k^P{}_1 \cdot S_1 \cdot R_1 \qquad \text{(Equation 1)}$$

$$S^P{}_2 = k^P{}_2 \cdot S_2 \cdot R_2 \qquad \text{(Equation 2)}$$

#### SMAD complex formation

Upon phosphorylation, two SMADs form complexes together with a third shared SMAD4 ($S_4$). We consider the formation of canonical complexes with either two $S_1$ proteins ($C_1$) or two $S_2$ proteins ($C_2$), with affinities of $k^C{}_1$ and $k^C{}_2$, respectively. The steady-state equations for SMAD complex formation are given by (Eq. 3-4).

$$C_1 = k^C{}_1 \cdot S^P{}_1 \cdot S^P{}_1 \cdot S_4 \qquad \text{(Equation 3)}$$

$$C_2 = k^C{}_2 \cdot S^P{}_2 \cdot S^P{}_2 \cdot S_4 \qquad \text{(Equation 4)}$$

#### Transcriptional response

The SMAD complexes translocate to the nucleus, where they act as transcription factors. To account for specific binding affinities, multiplicity of targets, and enzymatic activity of the complex, we consider each complex to have a target-specific transcriptional efficacy. Based on our dose-response data for the individual ligands, we find that each pathway only activates its corresponding reporter target. This allows us to conclude that the cross-transcriptional efficacies vanish for our transcriptional reporters. We thus consider a BMP target, B, that is activated only by the complex $C_1$ at a transcriptional efficacy of $e^B{}_1$ and a TGFβ target, T, that is activated only by the complex $C_2$ at a transcriptional efficacy of $e^T{}_2$. The steady-state equations for the transcriptional response are given by (Eq. 5-6).

$$B = e^B{}_1 \cdot C_1 \qquad \text{(Equation 5)}$$

$$T = e^T{}_2 \cdot C_2 \qquad \text{(Equation 6)}$$

#### Linearity of the transcriptional response

In this study, we use the assumption of linear transcriptional response (Equations 5,6). However, biological effects such as thresholds and cooperativity generally result in non-linearities. While these do not affect the overall nature of the crosstalk, they can contribute to qualitative effects. Specifically, using a simple non-linear dependence:

$$B = e^B{}_1 \cdot C_1 \cdot C_1 \qquad \text{(Equation 7)}$$

$$T = e^T{}_2 \cdot C_2 \cdot C_2 \qquad \text{(Equation 8)}$$

We find a more significant decrease of the BRE for smaller changes in TRE, and a more significant increase of the TRE for smaller changes in BRE (Figure S6A). This effect can also be observed in the experimental data.

#### Mass conservation equations

In addition to the steady state equations, we also consider the mass conservation equations for the SMAD proteins. These are given by (Eq. 9-11).

$$S_1 = S^0{}_1 - S^P{}_1 - 2 \cdot C_1 \qquad \text{(Equation 9)}$$

$$S_2 = S^0{}_2 - S^P{}_2 - 2 \cdot C_2 \qquad \text{(Equation 10)}$$

$$S_4 = S^0_4 - C_1 - C_2 \quad \text{(Equation 11)}$$

Together, this gives a set of 9 equations for 9 variables with 11 parameters.

#### A model with heteromeric complexes

The above model captures the known interactions within the intracellular part of the BMP and TGFβ pathway. We next extended our model to include the possibility of heteromeric SMAD complexes composed of a mixture of BMP and TGFβ-related SMADs. In the model, such a heteromeric complex, $C_{mix,}$ is composed of a single $S_1$, a single $S_2$, and $S_4$ with a binding affinity of $k^C_{mix}$. This complex controls the transcriptional response of the target genes B and T with efficacies of $e^B_{mix}$ and $e^T_{mix}$, respectively:

$$C_{mix} = k^C_{mix} \cdot S^P_1 \cdot S^P_2 \cdot S_4 \quad \text{(Equation 12)}$$

The resulting full model is described with 10 equations (Eq. 13-22) for 10 variables (Table S6) and contains 14 parameters (Table S7).

$$S^P_1 = R_1 \cdot k^P_1 \cdot S_1 \quad \text{(Equation 1)}$$

$$S^P_2 = R_2 \cdot k^P_2 \cdot S_2 \quad \text{(Equation 2)}$$

$$C_1 = k^C_1 \cdot S^P_1 \cdot S^P_1 \cdot S_4 \quad \text{(Equation 3)}$$

$$C_2 = k^C_2 \cdot S^P_2 \cdot S^P_2 \cdot S_4 \quad \text{(Equation 4)}$$

$$C_{mix} = k^C_{mix} \cdot S^P_1 \cdot S^P_2 \cdot S_4 \quad \text{(Equation 12)}$$

$$B = e^B_1 \cdot C_1 + e^B_{mix} \cdot C_{mix} \quad \text{(Equation 5)}$$

$$T = e^T_2 \cdot C_2 + e^T_{mix} \cdot C_{mix} \quad \text{(Equation 6)}$$

$$S_1 = S^0_1 - S^P_1 - ( 2 \cdot C_1 + C_{mix} ) \quad \text{(Equation 9)}$$

$$S_2 = S^0_2 - S^P_2 - ( 2 \cdot C_2 + C_{mix} ) \quad \text{(Equation 10)}$$

$$S_4 = S^0_4 - ( C_1 + C_{mix} + C_2 ) \quad \text{(Equation 11)}$$

#### Dimensional analysis

Next, we want to analyze the parameters of this model and determine the minimal set of parameters required to simulate the full spectrum of solutions. To do that, we use dimensional analysis, which allows us to reduce the number of meaningful parameters in a model. As the variables of the model are dimensionful, some changes in parameters only reflect a choice of units, and thus these parameters can be eliminated. In this model, we can describe all variables and parameters using 5 distinct dimensionful units - s, $r_1$, $r_2$, b, and t, as follows.

$[R_1] = r_1$

$[R_2] = r_2$

$[S_1] = [S^0{}_1] = [S^P{}_1] = [S_2] = [S^0{}_2] = [S^P{}_2] = [S_4] = [S^0{}_4] = s$

$[k^P{}_1] = (r_1)^{-1}$

$[k^P{}_2] = (r_2)^{-1}$

$[C_1] = [C_2] = [C_{mix}] = s$

$[k^C{}_1] = [k^C{}_2] = [k^C{}_{mix}] = s^{-2}$

$[B] = b$

$[T] = t$

$[e^B{}_1] = [e^B{}_{mix}] = b\ s^{-1}$

$[e^T{}_2] = [e^T{}_{mix}] = b\ s^{-1}$

$[B] = b$

$[T] = t$

Choosing specific natural units, we can fix some parameters. Without loss of generality, we can choose specific units for $R_1$, such that $k^P{}_1=1$, and for $R_2$, such that $k^P{}_2=1$. Similarly, by choosing appropriate units for B and T we can set $e^B{}_1 = e^T{}_2 = 1$. Finally, by rescaling the units of the SMAD molecules, we can set $k^C{}_2 = 1 / k^C{}_1$. We note that other choices for the units could equally be used, generating the same results, for example setting $k^C{}_1=1$ and keeping $k^C{}_2$ independent. However, our choice retains the symmetry of the equations (between BMP and TGFβ), and reflects the biological understanding that only the ratio between the affinities affects the behavior of the pathways. Overall, we use dimensional analysis to fix 5 of the parameters, resulting in a model with a 9-dimensional parameter space. We note that for the model without heteromeric complexes, a similar dimensional analysis results in a 6-dimensional parameter space.

#### Simulations

We systematically simulated the model across the parameter space and quantified the crosstalk. We analyzed the response on a 7-dimensional grid corresponding to the 9 model parameters excluding $R_1$ and $R_2$ which represents the input values for the simulations:

| | |
|---|---|
| $S_1$ | Logarithmically spaced between 0.01 and 100 |

| | |
|---|---|
| $S_2$ | Logarithmically spaced between 0.01 and 100 |
| $S_4$ | Logarithmically spaced between 0.01 and 100 |
| $k^C_1$ | Logarithmically spaced between 0.01 and 100 |
| $k^C_{mix}$ | Logarithmically spaced between 0.01 and 100 |
| $e^B_{mix}$ | Linearly spaced between 0 and 1 |
| $e^T_{mix}$ | Linearly spaced between 0 and 1 |

For each parameter set, we simulated the system for various levels of input signals $R_1$ and $R_2$. We note that we limit the overall efficacy of the heteromer complex ($e^T_{mix}$+$e^B_{mix}$) by the efficacy of a homodimeric complex ($e^B_1 = e^T_2 = 1$), so that an increase in the response will not be a simple artifact of an increase in efficacy. In particular, to measure the crosstalk on each pathway, we calculated the response when both inputs were high and compared with the response when only the corresponding receptors were high. To define a specific high level for each receptor, we simulated a dose-response curve for each individual receptor and used the level resulting in an activity of 90% of saturation. To generate the 2D response matrix, we used receptor levels activating the response at 95%, 90%, 75%, 50%, 25%, 10%, and 5% of saturation. We solve the resulting system of coupled reaction kinetics equations with the Python package Equilibrium Toolkit (EQTK)[84,85].
